# ARViS: A bleed-free multi-site automated injection robot for accurate, fast, and dense delivery of virus to mouse and marmoset brains

**DOI:** 10.1101/2024.01.15.575593

**Authors:** Shinosuke Nomura, Shin-Ichiro Terada, Teppei Ebina, Masato Uemura, Yoshito Masamizu, Kenichi Ohki, Masanori Matsuzaki

## Abstract

Genetically encoded fluorescent sensors continue to be developed and improved. If they could be expressed across multiple cortical areas in non-human primates, it would be possible to measure a variety of spatiotemporal dynamics of primate-specific cortical activity. Here, we develop an Automated Robotic Virus injection System (ARViS) for broad expression of a biosensor. ARViS consists of two technologies: image recognition of vasculature structures on the cortical surface to determine multiple injection sites without hitting them, and robotic control of micropipette insertion perpendicular to the cortical surface with 50-μm precision. In mouse cortex, ARViS sequentially injected virus solution into 100 sites over a duration of 100-minutes with a bleeding probability of only 0.1% per site. Furthermore, ARViS successfully achieved 266-site injections over the frontoparietal cortex of a common marmoset. We demonstrate one-photon and two-photon calcium imaging in the marmoset frontoparietal cortex, illustrating the effective expression of biosensors delivered by ARViS.

## Introduction

Genetically encoded calcium indicators (GECIs) have revolutionized the fields of neuroscience and functional biology^1–3^. Furthering their use, the recent development of genetically encoded neurotransmitter sensors has opened new opportunities to study the spatiotemporal dynamics of the release and spread of neurotransmitters such as glutamate and dopamine^4,5^. If these sensors could be expressed uniformly across multiple broad cortical areas in the non-human primate (NHP), it would be possible to measure the spatiotemporal dynamics of the inter-areal spreading of a variety of activities in motor, cognitive, and sensory processing that are well developed in the primate. In addition, the fluorescence intensity change in these sensors can be detected at the resolution of a single-neuron, and sometimes at a subcellular resolution.

To obtain cortex-wide imaging of fluorescence intensity changes in these sensors with a high signal-to-noise ratio, it is necessary that they are expressed broadly, uniformly, and strongly. The most common method for achieving this is the injection of adeno-associated virus (AAV) carrying the sensor gene into multiple sites. AAV-mediated GECI expression has been successfully demonstrated in macaque and common marmoset cortex^6–15^. Although a GECI-expressing transgenic marmoset was created, its expression level was very weak^16^; in general, gene expression in transgenic animals is weaker than high-titer virus-mediated gene expression. As an alternative, brain transfection of AAV variants such as AAV-PHP.eB and AAV.CAP-B10 through systemic transvenous administration is more promising^17^. However, transvenous transfection-mediated gene expression in the NHP brain is not sufficiently high for practical use of GECIs. Since transvenous transfection induces expression in other organs, such as liver, it might cause unintended malfunctioning of these organs, thereby limiting the viral dose and gene expression that can be obtained. Therefore, although molecular biologists are struggling to solve these problems, we need to adopt a method involving multiple local injections of the virus solution to image the sensor in NHPs.

Since the cortical area in which the gene should be expressed is broader in NHPs than in rodents, the number of injection sites and the total time for injections are greater in NHPs than in rodents. To make the injection procedure easier and more effective, automation using robotics could be a solution. To achieve this goal, two major problems need to be addressed. First, blood vessels should be avoided at the injection sites to prevent bleeding. Second, it is necessary to insert a glass micropipette filled with the virus solution into the target site perpendicular to the cortical surface to prevent slipping or bending of the pipette on the cortical surface and deformation of the cortical tissue. Although the technology for automated recognition of blood vessels has advanced^18–20^, it has rarely been used in real time immediately before surgery on the target tissue. Although micropipette insertion methods for in vivo patch-clamp recording have been automated, the insertion target is determined by the experimenter or by real-time imaging, and these methods do not have the ability to automatically avoid blood vessels^21–23^. For the purpose of deep brain stimulation therapy, CT- and MRI-assisted robotic technology has been developed for the automated insertion of electrodes into the brain while avoiding important intracranial structures such as nerve nuclei and ventricles^24^. Although this system is suitable for deep brain intervention, the misalignment range is approximately 1 mm from the target^25^. Therefore, insertion into the brain parenchyma is usually performed manually, and hemostatic treatment is necessary.

In this study, we developed an Automated Robotic Virus injection System (ARViS) that enables bleed-free multi-site automated injections for accurate, fast, and dense virus delivery into living brains. We solved the first problem of blood vessel avoidance by precisely measuring the three-dimensional (3D) structure of the cortical surface and applying deep learning-based image recognition of the vasculature structures with clustering-based optimization of multiple injection sites. We solved the second problem of perpendicular insertion by manipulating the micropipette with a six degrees-of-freedom robot under high-precision calibration. To demonstrate the imaging of ARViS-assisted expression of the GECI, we imaged different neuronal representations of orofacial and body movements over a 14 × 7 mm area of the frontoparietal cortex in an awake adult marmoset.

## Results

### Workflow of ARViS

ARViS consists of two technologies: robotic technology to precisely, smoothly, and sequentially insert a micropipette into multiple virus-injection sites in the cortex (Figure 1A–D), and image recognition technology to determine multiple virus-injection sites in the cerebral cortex without hitting blood vessels on the cortical surface (Figure 1E–I). The baseplate of a hexapod robot that had six degrees of freedom was fixed on the desk parallel to the vertical plane. The stage of the robot was attached to an injector connected to a micropipette, an infrared laser distance sensor to measure the 3D cortical surface shape, and a CMOS camera to image cortical blood vessels (Camera S) (Figure 1A). Anesthetized animals with the skull removed were set in a head-fixation apparatus on the base plate. The cortical surface and injection sites were determined in the XYZ coordinates in the robot base system frame {**B**} (Figure 1A, C, I), and the micropipette filled with solution was inserted into the cortical tissue at each site by a combination of XYZ movements, V rotation (along the Y axis), and W rotation (along the Z axis), under robotic control.

**Figure 1.**
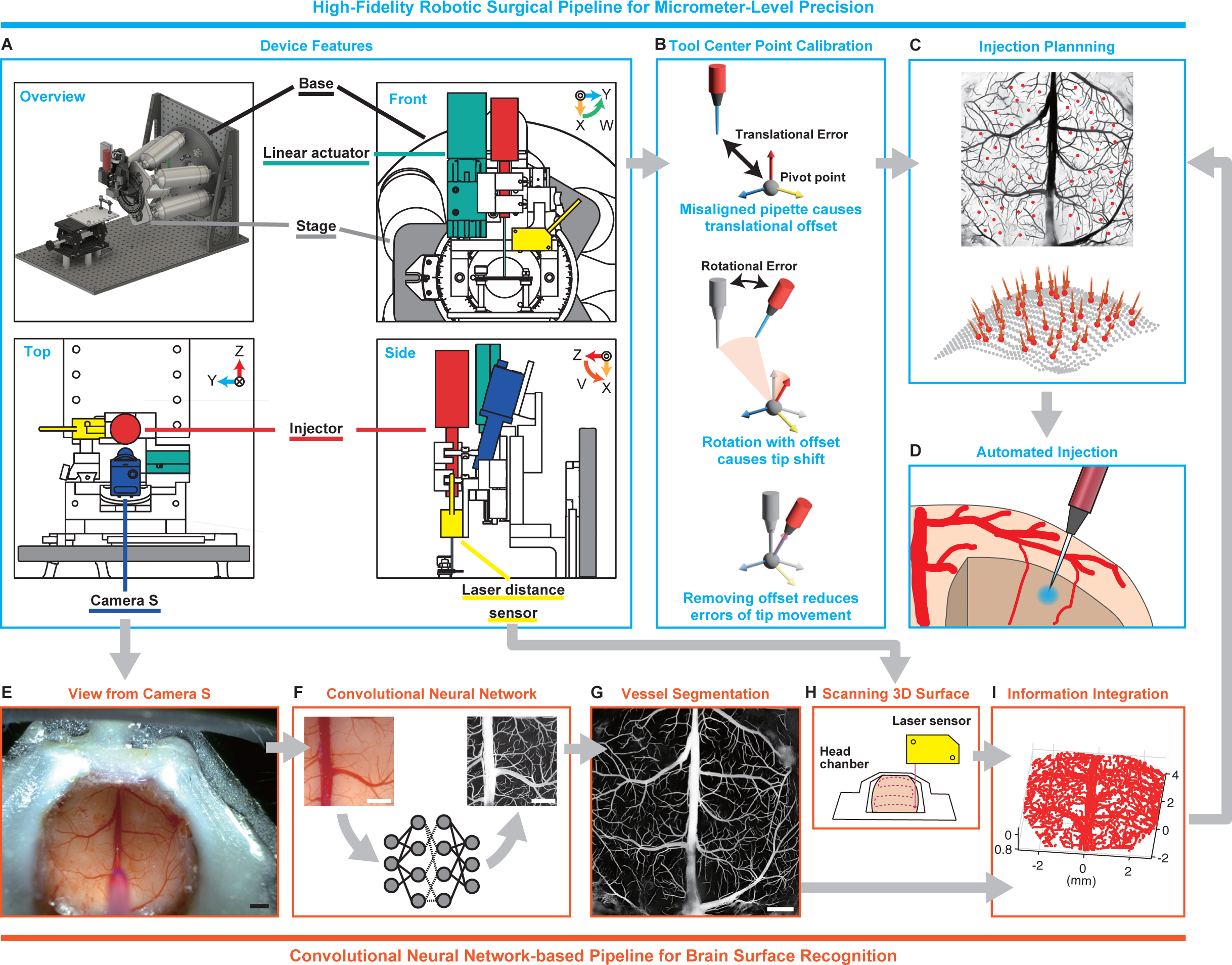
Workflow of an automated injection. (A) Schematic of the device features. An injector for connecting a glass micropipette for the virus injection (red), a linear actuator to move the injector up and down (green), a laser distance sensor for measuring the 3D cortical surface (yellow), and Camera S to capture the vasculature structure on the cortical surface (blue) were mounted on the stage of a six-axis hexapod robot. The animal’ s head was fixed in the apparatus on the base plate (the animal is not shown). Light orange, blue, and red arrows shown in the upper right of each inset represent the X, Y, and Z axes in the robot base system frame, respectively. The V (orange) and W (green) rotation directions are also shown. (B) Schematic of tool center point calibration. Top, any misalignment of the pipette tip position from the pivot point can cause an unintended translational offset from the desired position. Middle, when the pipette rotates around the pivot point, this offset can induce an unexpected shift in the position of the pipette tip. Bottom, minimizing this offset is essential for 3D manipulation of the micropipette. (C) A representation of multiple injection sites (red points) on the mouse cortical surface with vessel segmentation (black) and the directions of the pipette insertion (red lines in the bottom). The positions of the injection sites and the pipette angles for individual injections are designed based on the information from the pipeline for the cortical surface recognition (E–I). (D) Illustration of the automated injection of virus into the target point within the cortical tissue. (E) A representative image of the mouse cortical surface captured by Camera S shown in (A). Scale bar, 1 mm. (F) Schematic of the segmentation of blood vessels on the cortical surface with a convolutional neural network. The left image is an example of the input data, and the right image is its output data. Scale bars, 1 mm. (G) A representative stitched image of the vessel segmentation over the whole dorsal cortex of a mouse. Scale bar, 1 mm. (H) Schematic of 3D cortical surface scanning with the laser distance sensor. (I) An example of the projection of the vessel pattern onto the 3D cortical surface.

### Deep learning-based vessel segmentation

In the first step of the image recognition, the 3D cortical surface (an area over 8 × 8 mm centered on the bregma) of the anesthetized mouse was accurately scanned with the infrared sensor. On the basis of this surface scanning data, the distance between Camera S and the cortical surface was automatically adjusted and finely-focused surface images were acquired at 62 points over the neocortex (Figure S1A). Next, the blood vessels within a cropped area of each image (approximately 3.8 × 3.8 mm; Figure 1F, 2A and Figure S1B) were segmented using a deep learning-based convolutional neural network (CNN) named Spatial Attention-UNet (SA-UNet)^26^, which was originally designed for the segmentation of retinal fundus vasculature (Figure S2A). We initially trained the neural network using twenty images of the retinal fundus vasculature from the Digital Retinal Images for Vessel Extraction (DRIVE) dataset (Figure S2B, C). The trained model worked well for relatively narrow vessels, but not for thick vessels (Figure 2B), which might reflect the fact that the widths of the vessels on the mouse cortex range from less than 10 µm to more than 200 µm, thereby differing substantially from retinal vessels. Therefore, we re-trained the SA U-Net using an additional six images of cortical vessels in mouse brain, with vessel labels that were determined by the experimenter (Figure S2D, E). Although this addition improved the detection of thick vessels, some parts of them remained undetected (Figure 2B). To add more effective mouse cortical images to the training dataset, we adopted EquAL^27^, which calculates the difference in prediction between the original image and its left-right inverted image as the “consistency” of the inference, and used CNN-corrected segmentation that uses active learning^28^. This CNN-corrected segmentation reduced the time for the EquAL data labeling by approximately one-third (Figure S2F). The modified SA U-Net reliably detected thick vessels, even the thickest one in the midline of the cortex (Figure 1F, 2B and Figure S2G, H; see Methods for details).

**Figure 2.**
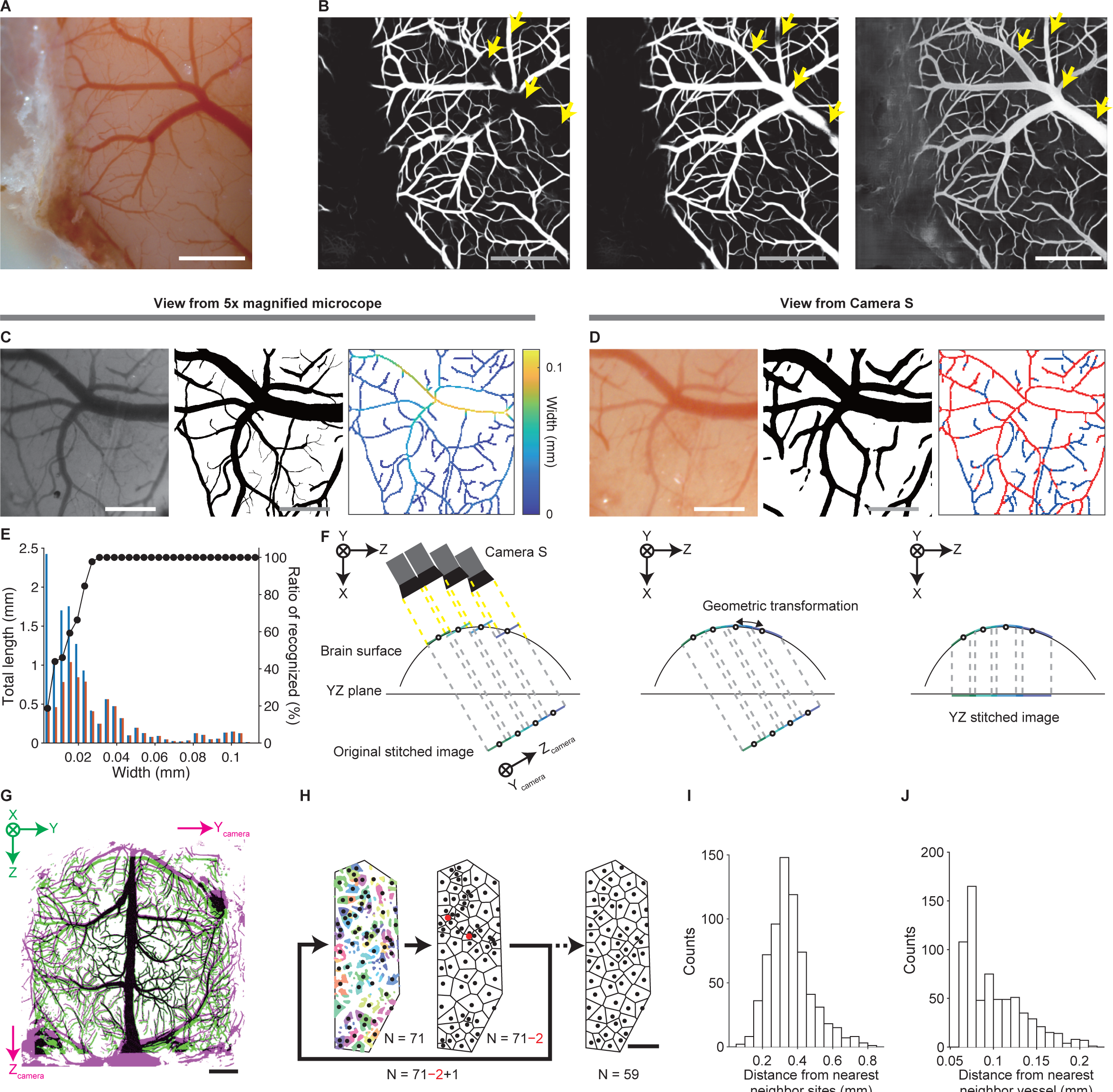
Vessel segmentation on the cortical surface. (A) An example input image of the mouse cortical surface for SA-UNet. Scale bar, 1 mm. (B) Example output images of vessel segmentation with SA-UNet when it was trained with only the DRIVE dataset (left), when it was trained with the DRIVE dataset and six manually created datasets (middle), and from the modified SA-UNet (right). The input image is the image shown in (a). Scale bar, 1 mm. Note that some parts of thick vessels were only segmented when the modified SA-UNet was used (arrows). (C) Left, an example image of the cortical surface captured by the microscope with a 5× magnification objective. Middle, manually created vessel segmentation of the left image. Right, skeletonized blood vessel mask of the middle image. The color of the skeleton corresponds to the width of the vessel at the corresponding point. Scale bar, 0.3 mm. (D) Left, an image captured by Camera S. The imaged area is the same as in (c). Middle, a vessel mask of the left image generated by the modified SA-UNet. Right, skeletonized blood vessel mask of the middle image. Vessels overlaid by the mask in (c) are shown in red, while the rest are in blue. Scale bar, 0.3 mm. (E) Histogram of the total lengths of the vessels that were detected on the 5× magnification image (blue) and those that were recognized by SA-UNet (red) for each bin width. A black plot for each bin width indicates the ratio of the latter length to the former length (n = 1 mouse). (F) Schematic of the transformation process from the original stitched image to the YZ stitched image. Left: Each image was acquired by adjusting the X position so that the position of the reference point where the pipette tip should be imaged matched the cortical surface, and these images were then stitched to form the original stitched image (Y_camera_-Z_camera_ plane). Circles on the cortical surface indicate the reference points and circles on the stitched images are their projection. Middle: By inferring the angle between the camera and laser optical axes, the original stitched image was mapped to the corresponding cortical surface with geometric transformation. Right: The image mapped on the cortical surface was projected to the YZ plane as the YZ stitched image. The camera angle and the curvature of the cortical surface are more emphatic than factual. (G) Representative overlay of the vessel segmentations of the original stitched image (magenta) and the YZ stitched image (green). Scale bar, 1 mm. (H) Diagram for the injection site planning algorithm. The initial number (N = 71 in this mouse) of sites (black dots) within the safety regions (colors) were determined according to the k-means clustering method. Different colors indicate different clusters. White areas indicate the safety margin region (with a safety margin distance of 65 μm in this mouse) and blood vessels. Next, two sites whose cluster areas including blood vessels were the smallest and second smallest (red dots) were removed and one new site was added. After these procedures were repeated approximately 10 times (eleven times in this mouse), the target injection sites were determined. Scale bar, 1 mm. (I) Histogram of the distance between the target injection site and the corresponding nearest neighbor site (n = 679 sites from seven mice). (J) Histogram of the distance between the target injection site and the corresponding nearest blood vessel (n = 679 sites from seven mice).

To determine the minimum size of the extracted blood vessels, we compared the model-segmented vessels with vessel structures that were manually determined from cortical images taken at a higher magnification under the microscope (Figure 2C, D). Vessels of ≥ 30 µm width were recognized by the model-based segmentation with 100% accuracy, whereas vessels with approximately 20 µm width were recognized with nearly 80% accuracy (Figure 2E). We assumed that if we avoided vessels with a width of approximately 20 µm or more, then serious bleeding would not occur when the injections were performed.

### Determination of multiple injection sites in the two-dimensional vessel segmentation image

The vessel segmentation images were stitched together to make one image covering the whole dorsal cortex (left in Figure 2F; original stitched image). However, the angle of the camera was tilted by approximately 15° against the optical axis of the laser (Figure 1A). This meant that the greater the angle between the optical axis of Camera S and the perpendicular line of the cortical surface, the more the imaged vessel thickness and position deviated. Thus, the original stitched image needed to be mapped onto the 3D cortical surface anew. The reference points on the 3D cortical surface that were estimated from the laser scanning were projected onto a two-dimensional (2D) image at a given angle and a geometric transformation was performed in this projected image to minimize the difference in the reference point positions between the transformed projected image and the original stitched image. Then, the angle with the minimal difference in the reference point positions was regarded as the angle between the optical axes of the laser distance sensor and Camera S. The vessel segmentation image in the original stitched image was then transformed to the image on the 3D surface shape according to this angle (middle in Figure 2F). When the 3D segmentation image was projected into the YZ plane to create a 2D image (YZ stitched image; right in Figure 2F), differences in vessel structure between the original stitched image and YZ stitched image were apparent around the areas where the curvature of the cortical surface was prominent (Figure 2G and Figure S1A). Thus, we considered that the image transformation procedure worked well.

Next, we used the YZ stitched image to determine multiple injection sites over the dorsal cortex. It was not possible to insert the pipette into each target injection site outside of the blood vessels without any error. To prevent it from hitting a blood vessel, we designated the cortical region that was a certain distance away from the blood vessels (safety margin distance) as the safety region. To distribute multiple injection sites as evenly as possible within the safety regions, these regions were divided into a number of clusters according to the k-means method^29^. The centroid of each cluster was chosen as a candidate injection site (Figure 2H). After this process was repeated approximately 10 times to remove the sites that were very close to their nearest neighbors, the target injection sites were determined. When the safety margin distance was set at approximately 65 μm and the number of final injection sites was 97 ± 11.8 (n = 7 mice), the distance from the nearest neighbor sites was 0.361 ± 0.046 mm and the distance from the nearest blood vessel was 0.100 ± 0.013 mm (n = 678 sites from seven mice) (Figure 2I). It took approximately 17 minutes to finish the laser scanning and take photographs of the cortical surface, perform vessel segmentation, create the YZ stitched image, and determine the target injection sites (Table S1).

### High-precision tool point calibration for 3D manipulation of the pipette tip with the hexapod robot

As described, it is necessary to control the robot to precisely translate and rotate the pipette tip to each target injection site on the cortical surface of the anesthetized animal. Since a pipette must be connected for every injection experiment, the position of the pipette tip should be calibrated for each experiment. The problem of how to find the coordinates of a tool tip added to a robot is generally referred to as the tool center point (TCP) calibration problem (Figure 1B). The traditional approach to the TCP calibration problem is to manually position the needle tip at various angles to align it with a mark^30^. However, this method is labor-intensive and lacks the precision needed for the current study. In addition, the use of fragile glass micropipettes in our study required a non-contact methodology. Therefore, for the initial step in the calibration we adopted a non-contact binocular-vision-based approach^31^.

First, we set two cameras (Camera R and Camera L) to capture the pipette tip from the side. Then, the three-dimensional position of the pipette tip in the binocular camera coordinate system determined by these two cameras was inferred (Figure 3A, B). Frame {**C**} is the binocular camera coordinate system frame. Next, the coordinates of the pipette tip in frame {**C**} are transformed to those in the robot base system frame {**B**} (Figure 3C). In the robot coordinate system, the “pivot point”, which serves as the effector point, can be set as the origin in the effector coordinate system (Figure 3C). Frame {**E**} is the effector frame. When the pipette tip and the pivot point are completely matched, the translation of the pivot point should move the pipette tip the same distance and the rotation around the pivot point should rotate the pipette tip by the same angle without any change in its position (Figure 1B). To make this match as precise as possible, the translation and rotation of the pipette tip under the control of the robot were repeated, its position at each movement end was measured with Camera L and Camera R, and then, the coordinate transformation matrix was determined to minimize the deviation of the pipette tip coordinates between frames {**C**} and {**B**} (see Methods for details).

**Figure 3.**
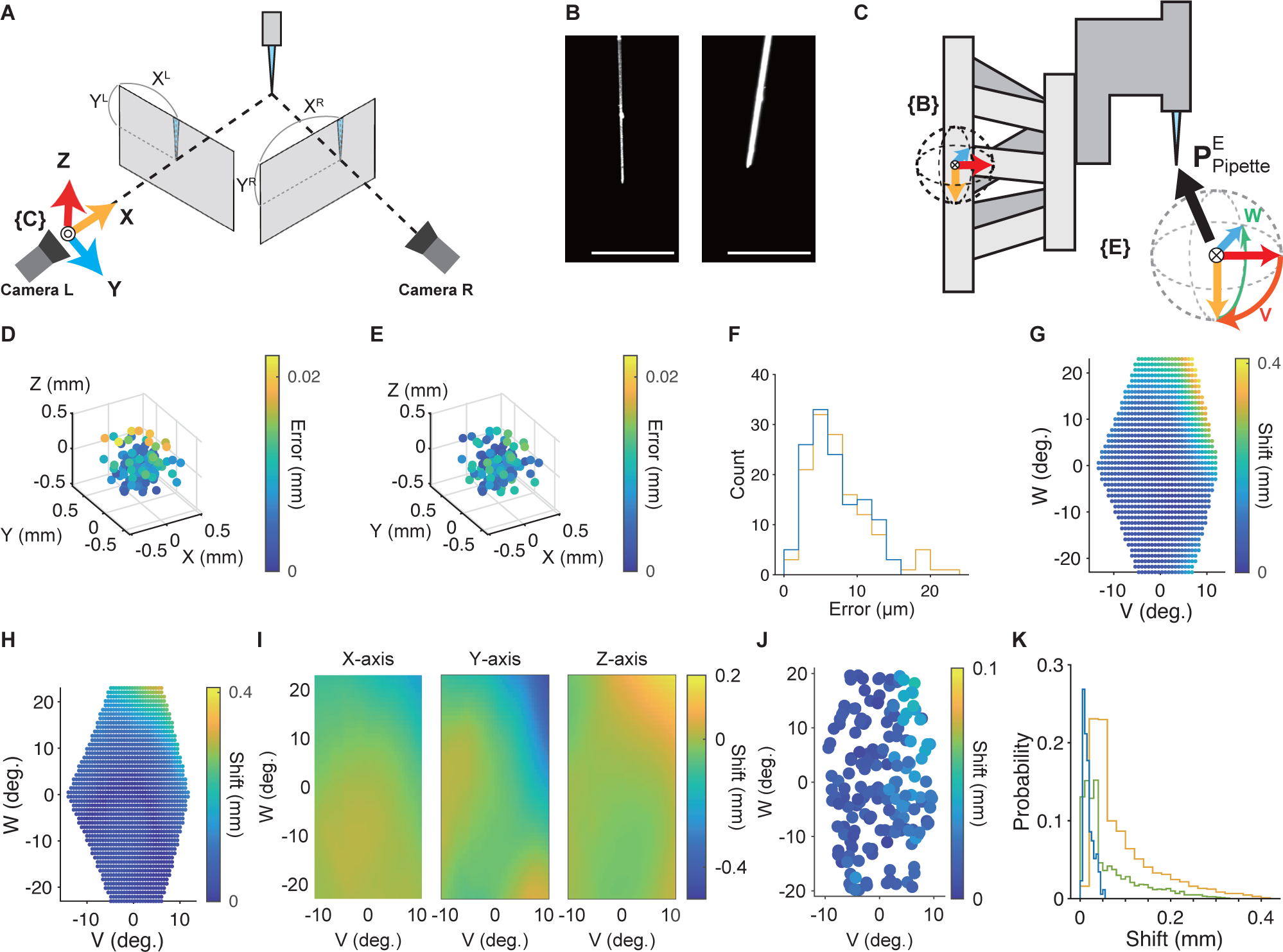
Tool center point (TCP) calibration. (A) Schematic of a binocular camera system for the 3D reconstruction of the pipette tip coordinates. 3D coordinates of the pipette tip are inferred from two 2D coordinates from the Camera L image (X^L^, Y^L^) and Camera R image (X^R^, Y^R^). (B) Example images of the glass micropipette captured by Camera L (left) and Camera R (right). The images were filtered, and the contrast was adjusted. Scale bar, 1 mm. (C) Coordinate system used for solving the tool center point calibration problem. **P**^E^_Pipette_ is a vector that indicates the initial position of the pipette tip in frame {**E**}. If **P**^E^_Pipette_ is accurately estimated, the pipette tip in frame {**B**} should be moved or rotated without any error. Light orange, X axis; cyan, Y axis; red, Z axis. The directions of V (orange) and W (green) rotations are also shown. (D, E) Representative plots of estimated pipette tip coordinates relative to the original tip coordinates before (D) and after (E) MSAC algorithm implementation. Only the coordinates regarded as inliers with the MSAC algorithm are shown for comparison (n = 131). The pseudo-color bar indicates the distance between the estimated pipette movement and the scheduled movement of the robot. (F) Histogram of the distance between the estimated pipette movement and scheduled movement of the robot without the MSAC algorithm (orange) and with the MSAC algorithm (blue). (G) A representative map of the pipette tip shift resulting from specific V and W rotation patterns before compensation for rotation error. The pipette tip was positioned at the coordinates predicted by the conventional TCP calibration algorithm (dots). The pseudo-color for each dot indicates the shift length. (H) A representative map of the pipette tip shift resulting from specific V and W rotation patterns after the first part of the compensation for rotation error. The pseudo-color for each dot indicates the shift length. (I) X- (left), Y- (middle), and Z- (right) axial maps of the pipette tip shift of the map shown in (H) after the interpolation using the thin-plate spline method. The pseudo-color indicates the length of the predicted shift. (J) A map of the pipette tip shift resulting from specific V and W rotation patterns after the compensation for rotation error. The pseudo-color for each dot indicates the shift length. (K) Histogram of the pipette tip shift for rotations with only conventional TCP calibration (orange), with grid searching after TCP calibration (green), and with all methods including the compensation (blue). Bin widths were determined using the Freedman-Diaconis method.

Although this calibration was able to bring the pipette tip close to the pivot point, some errors remained when translating or rotating the pipette tip. The first error was the error derived from the pipette detection on the captured images and camera calibration (Figure 3D). To more accurately estimate the coordinate transformation between frames {**C**} and {**B**} in the data with errors at many pipette-tip points, we implemented a robust estimation technique, the M-estimator Sample Consensus (MSAC) algorithm^32^. As a result, it was possible to estimate the pipette tip movement from the camera images with error of less than 15 μm for translational motion of 0.5 mm (Figure 3E, F).

The second error was caused by the robotic rotation. We noticed that the center point of the rotation could be shifted by a few hundred micrometers and that this shift depended on the rotation angle (Figure 3G and Figure S4A–D). To reduce the shift, we first used a voxel grid search to narrow down the deviation of the position of the pipette tip predicted from the pivot point (Figure S4E–H). As a result, the shift of the pipette tip in a rotation pattern (V of −7° and W of –15°) was reduced to less than 0.05 mm (Figure 3H). However, a maximum shift of 0.3 mm occurred with other rotation angles (Figure 3H). Therefore, we inferred the distribution of the shift in the V-W space by combining the grid research and regression model with a thin plate spline method^33^ (Figure 3I; see Methods for details). By compensating for the shift predicted by this model, 98.9% of the actual shift of the pipette tip position was reduced to less than 50 μm, and the average shift was 0.018 ± 0.001 mm (n = 175) (Figure 3J, K and Movie S1). When the XYZ position of the pipette tip was moved within a 3 × 8 × 8 mm space, which covered the mouse dorsal cortex (Figure S3A–C), and the rotation pattern was changed within a V range from −10° to 5° and a W range of ±20°, 99.8% of the shift caused by the rotation was less than 50 μm and the average shift was 0.021 ± 0.0002 mm (mean ± s.e.m., n = 1274) (Figure S4I, J).

It took 80–100 minutes to finish all processes of the TCP calibration (Table S1). All codes for the image recognition and robotic technology are listed in Table S1 and are publicly available at (https://github.com/nomurshin/ARViS_Automated_Robotic_Virus_injection_System/wiki).

### Validation of the precision of ARViS in the mouse neocortex

Next, we estimated the accuracy of the pipette insertion in the mouse cortex. We injected a fluorescent dye from the pipette tip into the mouse cortex and traced the pipette trajectory according to the fluorescence signal on post-fixed brain slices (Figure 4A). In this experiment, the target depth of insertion was set to 0.5 mm from the cortical surface. The injection depth was estimated at 0.50 ± 0.01 mm (mean ± s.e.m., n = 32 sites from one mouse) (Figure 4B) and the absolute misalignment was 0.05 ± 0.01 mm (Figure 4C). The lateral distance from each target injection site to the actual point of insertion was 0.038 ± 0.002 mm (n = 32 sites from one mice) (Figure 4E). These results show that even in the soft tissue of the cerebral cortex, a thin and bendable glass micropipette can be manipulated with a precision of approximately 50 μm.

**Figure 4.**
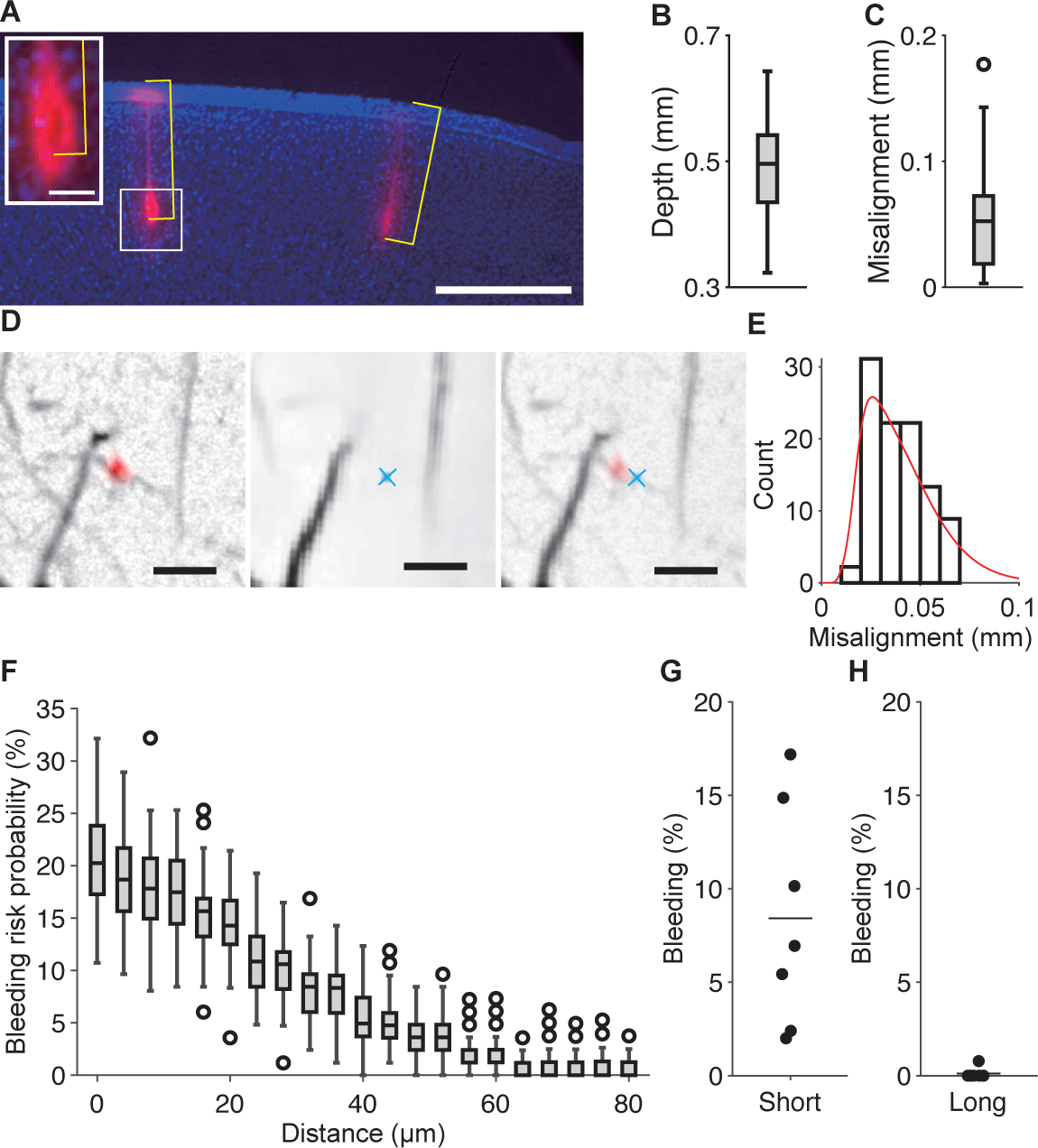
The accuracy of the pipette injection in vivo. (A) A representative coronal section of the post-fixed cerebral cortex of a mouse into which fluorescent dye (CM-DiI; red) was injected. The section was Nissl-stained (blue). The yellow lines indicate the depth of the injection from the cortical surface. When the putative pipette location was considered as a black void surrounded by the fluorescent dye, the deepest point of the void was measured as the depth of the injection (inset). Otherwise, the injection’ s depth was determined by measuring the deepest point of the fluorescent dye. Scale bar, 0.5 mm. Scale bar in the inset, 50 µm. (B) Box-and-whisker diagram of the measured depth (n = 32 sites from one mouse). (C) Box-and-whisker diagram of the absolute difference between the targeted and measured depths (n = 32 sites from one mouse). (D) Representative images showing the lateral misalignment of the pipette injection site. Left: A fluorescence (red, CM-DiI) image of the actual injection site superimposed on a bright field image. Middle: The target injection site (a blue cross) superimposed on a bright field image of the SA-UNet segmentation. Right: A composite image of the left and middle images. The lateral misalignment of the pipette tip was estimated as the distance between the center of the fluorescence spot and the target injection site. Scale bar, 100 μm. (E) Histogram of the lateral misalignment of the pipette tip (n = 32 sites from one mice). The red curve indicates a non-parametric approximation of the lateral misalignment distribution, calculated using a Gaussian kernel. (F) Box-and-whisker diagram of the bleeding risk probability against the safety margin distance (n = 100 simulations). (G, H) Rates of short (G) and long (H) bleeding when the pipette was injected perpendicular to the cortical surface at each site (n = 7 mice). The horizontal line represents the mean.

The risk of bleeding should depend on the distance between the target injection site and the nearest blood vessel. We therefore examined how the bleeding risk probability decreased as the safety margin distance increased. The bleeding risk probability was simulated as the proportion of injection sites corresponding to a blood vessel structure. Based on the distribution of the measured misalignments (shown as red plots in Figure 4E), the displacement was probabilistically added to 120 target injection sites that were determined when the safety margin distance was set in one cortical image. Then, the total number of virtual hits of the vessel was counted as the bleeding risk probability. The bleeding risk probability was near zero when the safety margin distance was more than 60 µm (e.g., 0.85 ± 0.09% at 64 µm) (Figure 4F). Therefore, we set the safety margin distance to 65 µm.

### Multiple injections in the mouse dorsal cortex without bleeding

The robot moved the pipette to the 3D coordinates of a target site along the perpendicular line at that site and inserted the tip into the cortical tissue. After injecting the fluid, the pipette was pulled back and moved into position above the next target injection site. This procedure was performed continuously for all sites in each animal. It took approximately 1 minute to complete the series of movements at one injection site.

To determine the frequency of actual bleeding occurrence in the mouse cortex after automatic multiple injections, we conducted a total of 679 injections in seven mice (AAV solution in six mice and the dye solution in one mouse). In these mice, the range of the insertion angles was within ±10° in the V rotation and ±20° in the W rotation (Figure S5A, B). In 8.43% ± 2.24% of the injection sites, short-duration bleeding occurred, with this defined as bleeding that stopped within 30 s after starting (Figure 4G). However, this short bleeding did not apparently affect the cortical structure after it had disappeared. By contrast, long bleeding, which was defined as bleeding lasting for more than 30 s, caused brain swelling or changed the vascular structure on the cortical surface and needed some hemostatic treatment. Long bleeding occurred in only one site in the seven mice (0.11% ± 0.11% of the injection sites) (Figure 4H). Thus, we concluded that the estimation of the blood vessel structure on the 3D cortical surface, determination of the injection sites in the 2D vessel segmentation image, and 3D manipulation of the glass micropipette worked very well.

### Multipoint injections into the marmoset frontoparietal cortex without bleeding

Once we were able to demonstrate the effectiveness of multi-site injections in mice using ARViS, we applied it to wide-field calcium imaging of the adult marmoset cortex. To enhance GECI expression by a tetracycline-inducible gene expression system^7^, a mixture of two AAVs encoding hsyn-tTA and TRE-tandem-GcaMP6s^8,34^ was injected into a total of 266 sites over the right frontoparietal cortex (16 × 8 mm) in one marmoset (Figure 5A). The range of the insertion angles was –20° to 0° in the V rotation and –10° to +20° in the W rotation (Figure S5C, D). The injected cortical areas were roughly inferred from intracortical microstimulation (ICMS) and a stereotaxic atlas^8,35,36^ as the area from the rostral part of the dorsal premotor cortex (6DR) to areas PE and PFG of the parietal cortex (Figure S6A). The entire injection process was divided into three cycles and the calibration was performed for each cycle because of changes in the surface shape over the surgery period and the limited volume of solution that could be loaded into the glass micropipette (Figure 5A). Cycles 1, 2, and 3 consisted of 89, 87, and 90 injections, respectively, at a depth of 0.5 mm from the cortical surface. Short bleeding was observed at seven sites (2.6%) in total (zero in cycle 1, two in cycle 2, and five in cycle 3), but no long bleeding was observed. Thus, ARViS was also effective in the marmoset cortex.

**Figure 5.**
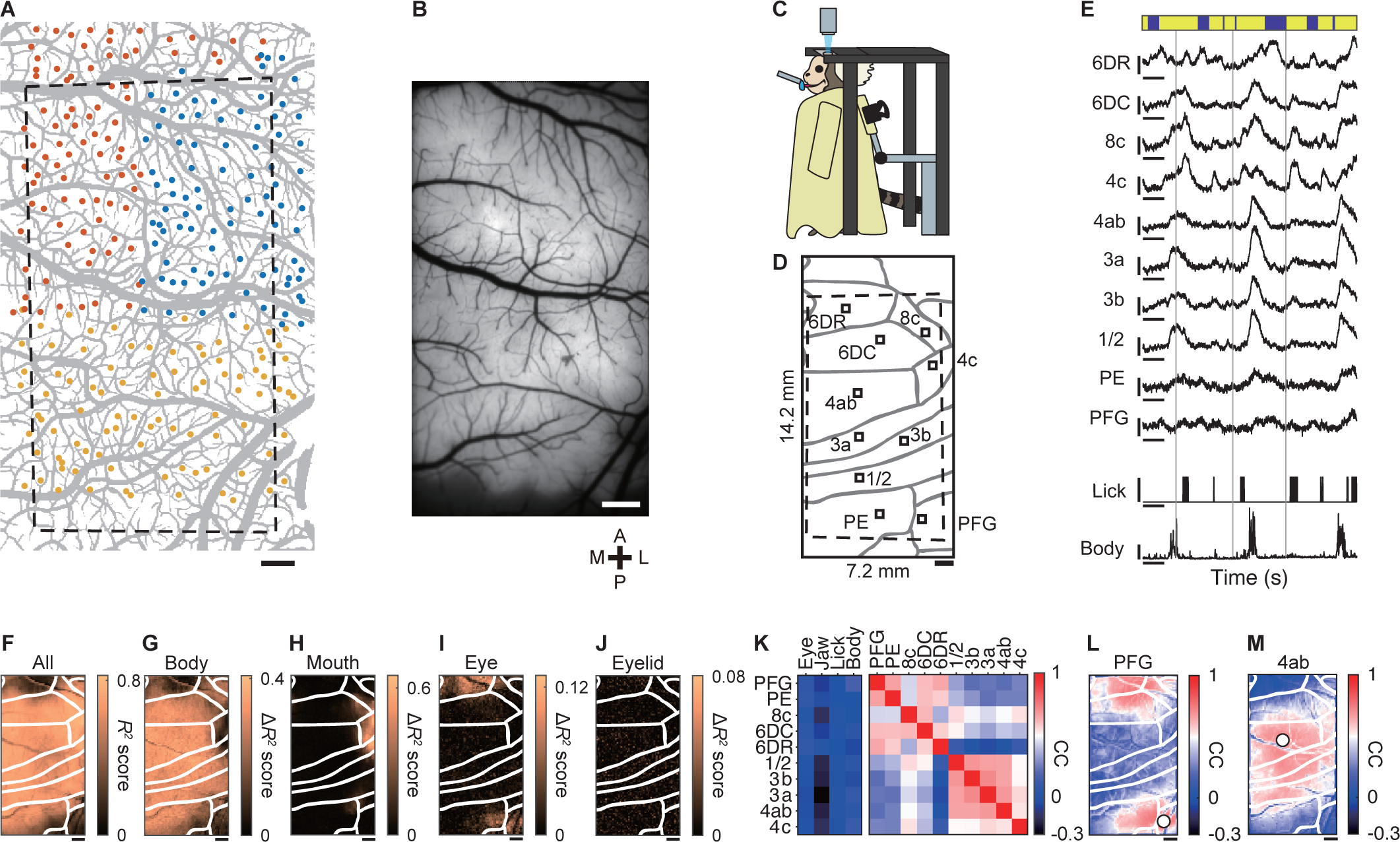
Wide-field calcium imaging of the frontoparietal cortex in an awake marmoset. (A) A vascular image of the marmoset frontoparietal cortex that was segmented by the modified SA-UNet (in gray) with target injection sites (colored dots). Blue, red, and yellow dots represent the first, second, and third sections, respectively. The black dashed frame represents the field of view for the fluorescence microscopy. The slightly distorted shape was caused by the affine transformation of the frame shown in (B), which was based on the blood vessel structure. This might be due to the elapsed time from the injection and distortion of the cortex caused by installation of the glass window. Scale bar, 1 mm. (B) Fluorescence image of the same area as (A) 7 weeks after the virus injection. Scale bar, 1 mm. A, P, M, and L stand for anterior, posterior, medial, and lateral, respectively. (C) Schematic of the experimental setup for wide-field one-photon calcium imaging of the marmoset in an awake head-fixed condition. (D) Putative cortical areas within the imaging window (black frame). The dashed frame indicates the field of view (14.2 × 7.2 mm). Parcellation is based on the stereotaxic atlas and ICMS (see Figure S6A). Small black squares represent ROIs for the corresponding cortical areas. Scale bar, 1 mm. (E) Representative time courses of Δ*F/F* at the ROIs of ten cortical areas shown in (C), licking, and body movement. The vertical gray lines show the reward timing. The vertical scale bars on the fluorescence traces represent 0.1 Δ*F/F*. The vertical scale bars with a lick represent a binary value of 1 and those with normalized body movement represent an intensity value of 1. The horizontal scale bars represent 5 s. The blue bars at the top indicate the quiet periods, whereas the yellow bars at the top indicate the periods outside of the quiet periods. (F) Spatial map of the accuracy (*R*^2^) of the encoding model for predicting the neural activity with all behavioral variables. The pseudo-colors represent *R*^2^ values. Scale bar for (F–J), 1 mm. (G–J) Spatial maps of the unique contribution (Δ*R*^2^) of the body (G), mouth (H), eye (I), and eyelid (J) variables to the explanation of the neural activity. The pseudo-colored bar represents the Δ*R*^2^ score. (K) Correlation matrix during the quiet periods between the cortical areas and movement variables (left), and between cortical areas (right). Two clusters between cortical areas were determined using hierarchical clustering. The pseudo-colored bar represents the correlation coefficient (CC). Note that although the eye movement was not considered in the definition of the quiet periods, all neuronal activity showed very weak correlations with eye movement. (L, M) Correlation maps between seed pixels (indicated by white filled circles) located in area PFG (L) and area 4ab (M) with the other pixels. The pseudo-colored bar represents the correlation coefficient (CC).

### Behavior-related cortical activities revealed by wide-field one-photon calcium imaging of an awake marmoset

Wide-field epi-fluorescent imaging successfully showed uniform fluorescence expression over the area of 14 × 7 mm at 7 weeks after the injections (Figure 5B). Then, we conducted wide-field one-photon calcium imaging in an awake head-fixed condition^8,9^ over three sessions (days) (Figure 5C). Licking frequently occurred following the water delivery from a spout in front of the animal’s mouth, whereas other body movements also occurred apart from the licking timing. When the neuronal activity (relative fluorescence change Δ*F*/*F*) in regions of interest (ROIs) in 10 cortical areas was extracted, the somatosensory-primary motor areas (1/2, 3a, 3b, and 4ab) appeared to increase activity when body movements occurred, and the ventral areas 4c and 8c appeared to increase the activity when licking occurred (Figure 5D, E). The time course of Δ*F*/*F* clearly shows that area 4c activity was related to licking and area 4ab activity was related to body movements (Figure S6B–E).

To reveal how the behaviors were coded over the imaging field, we constructed linear ridge regression models with behavioral variables to reconstruct the fluorescence signals in each pixel^37,38^. To assess the models’ effectiveness in capturing neural activity across various cortical areas, we calculated the 10-fold cross-validated coefficient of determination (denoted as cv*R*^2^)^37,38^. The model accounted for 54.9% ± 2.5% (mean ± s.e.m., n = 3 sessions) of the neuronal activity (Figure 5F). Subsequently, we estimated the unique contribution (Δ*R*^2^) of each variable^37^ (see Methods for details). Body movement was widely represented, with areas 3 and 4 at the center (Figure 5G). Mouth movement contributed strongly to the activity of area 4c (Figure 5H). Eye movement contributed to the activity of parts of area 6DR, area 8c, and a region near the boundary between areas PE and PFG, but eyelid movement was not strongly represented in any area (Figure 5I, J). Area 4c is responsible for facial movement, area 4ab is responsible for forelimb and body movements, and area 8c and the intraparietal areas are responsible for eye movement^36,39–41^ Thus, wide-field one-photon calcium imaging clearly showed the different representations of different movements in the marmoset frontoparietal cortex.

### Wide-field calcium imaging reveals distinct frontoparietal networks

To assess the functional connectivity that was not directly related to movements, we extracted the neuronal activity during the quiet state, which we defined as the time period excluding that from 0.3 s before to 2 s after the occurrence of licking or body movement. Then, we calculated correlations in the fluorescence change between the areas during this quiet state. A hierarchical clustering method was used to group the areas into two clusters: one included most of the somatosensory-primary motor areas, and the other encompassed the regions anterior and posterior to those in the first cluster (Figure 5K). The spatial patterns of these correlations were also visualized in a seed correlation map with areas PFG and 4ab serving as the seeds (Figure 5L, M). Anatomically, area 6DR receives axons from lateral and caudal parietal areas (e.g., area PFG and the intraparietal area)^40^. Thus, the correlation structure revealed by the linear regression analysis should reflect the anatomical connectivity, although the cortical structural map was only roughly inferred. These results indicate that ARViS-assisted wide-field calcium imaging showcased correlation patterns over the marmoset frontoparietal cortex.

### Behavior-related neurons revealed by two-photon calcium imaging of the marmoset motor cortex

Finally, we conducted two-photon calcium imaging of a field in area 4c (field 1) and a field in area 4ab (field 2) in the awake condition (Figure 6A–F). Each field was imaged at different depths on different days. We detected 143 and 159 active neurons in fields 1 and 2, respectively. Consistent with the results from one-photon imaging, the neurons in field 1 responded more strongly to the water delivery than the neurons in field 2 (Figure 6G–I). The licking was decoded much better by individual neurons in field 1 than in field 2, whereas the body movement was decoded better by those in field 2 than in field 1 (Figure 6J, K). The difference in decoding prediction accuracy was also prominent when the activities of individual neurons were summed over each field as an estimate of the population activity (Figure 6L, M). These findings suggest that the behavior-related population activity revealed by wide-field one-photon calcium imaging well reflected the average behavior-related activity of individual neurons. ARViS-assisted broad gene expression allowed us to reveal both the neuronal-averaged activity and the activity of multiple individual neurons in the marmoset cortex.

**Figure 6.**
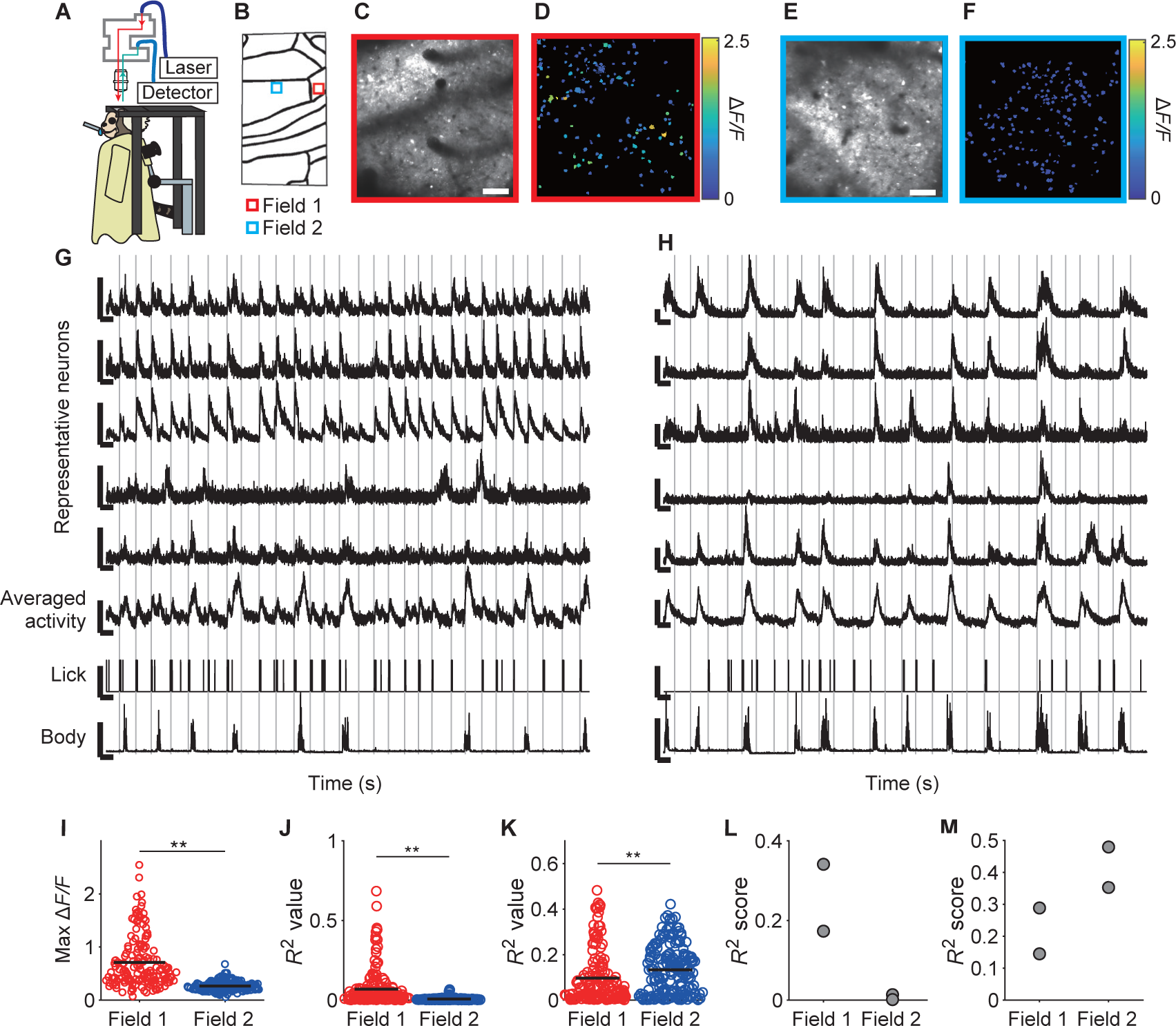
Two-photon calcium imaging of movement-related neurons in the marmoset motor cortex. (A) Schematic of the experimental setup to perform two-photon calcium imaging in the awake condition. The microscope body was tilted so that the excitation laser was incident perpendicular to the cortical surface. (B) Locations of two fields in which two-photon imaging was conducted. (C, E) Time-averaged and contrast-adjusted two-photon images for field 1 (C) and field 2 (E). Scale bars, 100 µm. (D, F) Neuronal somata extracted by the CNMF algorithm (Pnevmatikakis et al., 2016) from field 1 (D) and field 2 (F). Each neuronal soma is colored according to the maximum value from the trial-averaged Δ*F/F* trace during the period spanning 1 s before to 8 s after the reward timing. (G, H) Representative traces of five neurons from field 1 (G) and field 2 (H), averaged activity of all active neurons in each field, lick frequency, and body movement. The vertical gray lines show the reward timing. The vertical scale bars on the fluorescence traces represent 500% Δ*F/F*. The vertical scale bar with averaged activity represents 100% Δ*F/F*. The vertical scale bars with lick represent an intensity value of 1 (a.u.). The vertical scale bars with movement represent an intensity value of 5 (pixel). The horizontal scale bars represent 10 s. (I) Maximum values of trial-averaged Δ*F/F* traces for a period spanning 1 s before to 8 s after the reward timing in field 1 (red, n = 159 from 2 sessions) and field 2 (blue, n = 182 from 2 sessions). Horizontal bars indicate the means. **p = 5.0 × 10–23, Welch’ s test. (J, K) The prediction accuracy (*R*^2^) of individual neurons in fields 1 (red, n = 159 from 2 sessions) and 2 (blue, n = 182 from 2 sessions) for the licking frequency (J) and body movement (K). Horizontal bars indicate the means. **p = 2.6 × 10–10 (J) and 0.0028 (K), Welch’s test. (L, M) The prediction accuracy (*R*^2^) of the averaged activity in fields 1 (n = 2 sessions) and 2 (n = 2 sessions) for the lick frequency (L) and body movement (M).

## Discussion

In the current study, we developed an automated injection system capable of performing sequential multipoint injections into the dorsal cortex of mice and a marmoset. The injection point accuracy was 50 μm along the horizontal and perpendicular axes. The probability of long bleeding requiring hemostatic treatment was only 0.1% per injection. Furthermore, wide-field calcium imaging in a marmoset showed that this system allowed the broad expression of sensor proteins.

Although the technology for blood vessel recognition is advanced, especially in fundus examination, image recognition in fundus examination and robotic surgery have rarely been performed consecutively. A neurosurgical instrument that uses deep learning-based segmentation to automatically recognize blood vessels in the brain during robot-assisted surgery and suppress high-risk movements has been designed^42^, and while it showed good accuracy in a clay-based phantom study, it was not tested in in vivo living tissue. For this neurosurgical instrument, the distance between the target and vessel was set to 2 mm, which is much longer than the distance in our study (65 μm). Therefore, to our knowledge, the present study is the first to apply vessel recognition technology to biological intervention with high precision. In the current study, we segmented blood vessels only from the cortical surface, although there are also many vessels within the cortical tissue. However, cortical vessels tend to run perpendicular to the surface^43^, and thus many of them exist beneath the vasculature structure on the cortical surface, and they might therefore be well separated from the pathway of pipette insertion and the bleeding probability might be very small.

In the field of neurosurgery, the ROSA® system is a representative example of a robotic platform that was developed to automate the insertion of electrodes for deep brain stimulation therapy. By superimposing CT and MRI images on the patient’s head using a metal frame attached to the patient’s skull as a reference point, the system allows electrodes to be inserted while avoiding nerve nuclei and ventricles^24^. Although this system is suitable for deep brain intervention, it has an error of approximately 0.8 mm from the target^25^, which is not sufficiently accurate for virus injection into rodents and NHPs. In other robotic systems, such as NeuroMate and iSYS1^25,44–46^, CT images are used for registration. The diameter of the electrodes used for deep nuclei stimulation is more than 1 mm^47^, which is much longer than the diameter of the micropipette tip (0.03 mm). Therefore, these electrode insertion systems require hemostatic treatment during insertion into the brain parenchyma. The present study solved this problem by performing high-precision surface measurement using a laser distance sensor and high-precision calibration using cameras to accurately position the needle tip with an error of less than 50 µm. Furthermore, we demonstrated that even in soft cortical tissue with a curved surface, the misalignment was still approximately 50 µm along the axes parallel and perpendicular to the cortical surface.

Needle interventions in the brain are very common in neuroscience, and include single electrode recording and stimulation, multi-electrode recording^48,49^, and patch-sequencing, a combination of whole-cell patch-clamp recording and RNA sequencing^50^. The techniques presented in this study could be incorporated into the automated large-scale implementation of these methods. Even in deep brain stimulation, some of the algorithms developed in the current study may be applicable to CT angiographic images or magnetic resonance angiography to facilitate automatic determination of the best no-bleeding penetration pathway for the electrode and accurate manipulation of the electrode to the target area. The real-time control of image recognition and robotic technologies in ARViS would potentially be beneficial in many research and clinical medicine fields.

Apart from multiple microinjections, there are other methods for transfecting AAVs into NHP brains. For example, in-utero virus injections may be possible. However, developmental effects on the transfected gene are inevitable; it takes a year to obtain a young adult, even in the marmoset, and it is not clear whether a sufficiently large number of neurons will strongly express the sensors.

Although transvenous transfection of AAV variants such as AAV-PHP.eB and AAV.CAP-B10 will undoubtedly be further improved, even in NHPs^17^, the effectiveness of the transfection and the gene expression level in cortical neurons is unlikely to be sufficiently high for functional imaging in adult NHPs^51,52^. Thus, the systematic bleed-free multiple microinjection provided by ARViS should be highly useful for cortex-wide imaging of a variety of sensors in adult NHPs.

In contrast to NHPs, we now discuss whether ARViS is useful in research using mice that have many transgenic lines and Cre-lines. Relatively uniform expression or cell type-specific expression of sensors is possible by crossbreeding the appropriate Cre-line mice and sensor gene-expressing flox-lines. Even if there are no new sensor gene-expressing lines, they can be created within a few months. However, it takes more time to crossbreed multiple lines with different color sensors. By contrast, transvenous infection with AAV-PHP.eB is easy and quick, different sensors can be simultaneously transfected, and the sensors are widely expressed over the whole brain.

However, a problem with this approach is that it is not possible to limit gene expression to areas that are not genetically defined. An advantage of ARViS is its ability to realize spatially-selective gene expression independent of endogenous expression patterns and anatomical boundaries in both rodent and NHP cerebral cortices. ARViS promises a cross-species approach to many questions related to a variety of functions of the cerebral cortex, questions that have been difficult to solve with previous methods alone.

## Limitations of this study

The limitations of ARViS include the long calibration time (90 minutes) for the glass pipette position and the limited volume of solution (∼4 µL) for each pipette. The currently used injector limits the amount of virus that can be loaded. This problem can be overcome using a syringe with a large capacity as an injector. In addition, the current system was unable to compensate for deformation of the surface landscape after the AAV injection started. If cerebral edema is caused by vessel damage, it induces further bleeding and changes the cortical surface structure. This limitation could be overcome by periodic scanning of the surface during the injection process or by checking the position of the target injection site immediately before inserting the pipette. However, these steps would increase the time required. The ARViS system should be further improved with consideration of the trade-off between accuracy and speed.

## Supplementary information

The supplementary information includes six figures, one table, and one movie.

## Acknowledgments

We thank M. Nishiyama, Y. Hirayama, M. Hirokawa, and Y. Takahashi for animal handling, and Y. Watakabe, M. Takaji, and T. Yamamori for providing AAV constructs. This work was supported by Grants-in-Aid for Scientific Research on Innovative Areas (17H06309 to M.M., 19H05307 to T.E., and 21H00302 to T.E.), for Transformative Research Areas (A) (22H05160 to M.M. and 23H04977 to T.E.), for Scientific Research (A) (19H01037 and 23H00388 to M.M.), for Scientific Research (B) (20H03546 to T.E.), and for Early-Career Scientists (20K15927 and 23K14284 to S.-I.T) from the Ministry of Education, Culture, Sports, Science, and Technology, Japan; AMED (JP19dm0207069 to M.M.; JP15dm0207001 to M.M.; JP18dm0207027 to M.M.; JP19dm0207085 to T.E. and M.M.); and the Tokyo Society of Medical Sciences (to S.-I.T. and T.E.). This work was also supported by the program for Brain Mapping by Integrated Neurotechnologies for Disease Studies (Brain/MINDS) from AMED under Grant number JP21dm0207111.

## Author Contributions

S.N., S.-I.T., and M.M. designed the experiments. S.N. and S.-I.T. constructed the robotic system and conducted mouse experiments. S.N. performed most of the programming, optimized and manipulated the robot system, and analyzed the data. T.E. conducted surgery and imaging for the marmoset experiments. M.U., Y.M., and K.O. designed and prepared AAVs. S.N., S.-I.T., and M.M. wrote the paper, with comments from all authors.

## Methods

### Robot construction

The base of a six-axis hexapod robot with a maximum travel range of 100 mm in translation and 60° in rotation (Hexapod H-820; Physik Instrumente, Karlsruhe, Germany) was fixed vertically onto the base plate, and an injector (Nanoject III; Drummond Scientific, Broomall, USA), a laser distance sensor (HG-C1050-P; Panasonic, Osaka, Japan), and a CMOS camera (acA4024-29um; Basler, Germany) (Camera S) were attached to the stage of the robot via the original holding fixture. The original holding fixture was first created with a 3D printer but was then replaced with a metal one. The injector was attached to a programmable linear actuator (MTS50/M-Z8; Thorlabs, Newton, USA) that was connected to a computer via a USB port and could be moved backward during surface scanning. The injector and the laser distance sensor were connected to the Hexapod controller through analog signal cables, allowing them to communicate with the computer via the Hexapod. The camera (referred to as Camera S) was fixed approximately 15° relative to the vertical axis to capture the cortical surface (Figure 1E) and the tip of the glass micropipette. During the step of the calibration of the micropipette tip, two additional cameras (acA1440-220um; Basler, Germany) were set to detect it on the surgical plate via attachments. All these cameras were connected to the computer via USB ports. The code for the ARViS program was written and run in MATLAB (R2020a, R2021b; MathWorks, US).

### Tool center point calibration problem

Calculating the position of the pipette tip is equivalent to solving a tool center point (TCP) calibration problem. Normally, this problem is solved by bringing the needle tip into contact with a specific point and rotating the needle while maintaining that contact^30^. However, since the micropipettes used in this study were very fragile, a non-contact method was required. Zhang et al.^31^ proposed a binocular vision-based approach to this problem. Therefore, we adapted this method as: “1. Inference of the 3D coordinates of the pipette tip with the binocular camera system” and “2. Transformation of the 3D pipette coordinates to the frame centered at the pivot point”. Furthermore, since the AAV injection into the neocortex required a higher level of precision than that of Zhang et al., we developed two calibration methods to minimize errors that were not considered in the previous research: “3. Compensation for translation error” and “4. Compensation for rotational error”.

### 1. Inference of the 3D coordinates of the pipette tip with the binocular camera system

#### Pipette tip detection

A glass micropipette filled with the desired solution was connected to the injector. With the actuator, the position of the pipette tip was fixed to be at the focal plane of Camera S. In the images acquired by the two other cameras (Camera L and Camera R) for taking pictures of the pipette tip from the side, the origin of the two-dimensional coordinate frame is located at the top left corner (Figure 3A). The coordinates of the pipette tip within this frame were obtained as follows. First, the positions of the two cameras were fine-tuned so that the pipette tip could be clearly captured. Second, the background gradation in illuminance was removed through applying a top hat filter with a square structuring element to the captured images. Then, the images were binarized by thresholding them according to intensity values (Figure 3B). Finally, those binarized pixels with a value of one and the top five highest y coordinates were detected, and the x and y coordinates of these pixels were averaged.

#### Camera calibration

The two-dimensional frames on the captured images need to be transformed to a three-dimensional camera coordinate system. Frame {**C**} is the camera coordinate system frame. We aimed to estimate the camera matrices ***C***^*L*^ and ***C***^*R*^, which project the three-dimensional coordinates of the pipette tip 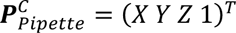 in the frame {**C**} centered at Camera L into the two-dimensional coordinates of the pipette tip 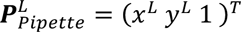 and 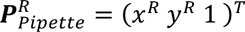, which were captured by Camera L and Camera R, respectively.

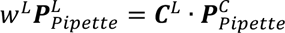

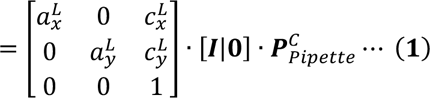

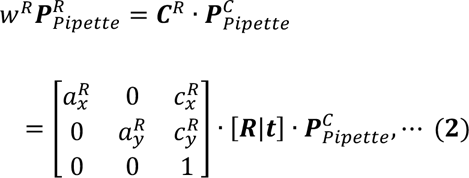

where

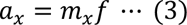

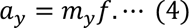

Values with a superscript L or R indicate the values for Camera L or Camera R, respectively; *f* represents the focal length of the camera (mm); and *m*_*x*_ *and m*_*y*_ are the scales in x and y directions (pixel/mm), respectively. (*c*_*x*_, *c*_*y*_) represent the optical center in the images (pixel). These parameters are intrinsic parameters of the cameras. ***R*** is a 3 × 3 rotation matrix relative to Camera L, and ***t*** is a 3 × 1 translation vector from Camera R to Camera L. These rotations and translations are extrinsic parameters of the cameras. ***C*** is a camera matrix that represents the transformation, which is composed of intrinsic and extrinsic parameters. Camera L needs neither translation nor rotation to project to Camera L itself; thus, the extrinsic parameters are expressed as [***I***|**0**], where *I* is an identity matrix. For the camera calibration, the inference of both intrinsic parameters and extrinsic parameters is necessary. The intrinsic parameters of the camera were estimated at the first use because they were unique to each camera and were generally invariant in daily use. For the extrinsic parameters, the position and angles were changed to allow focus on the pipette tip, so they were therefore estimated every time the pipette was replaced.

The intrinsic parameters were calculated by correlating grid coordinates obtained by moving the pipette tip in 0.2 mm intervals using the robot. This procedure was performed using the “estimateCameraParameters” function in Matlab’s Computer Vision Toolbox. The extrinsic parameters, the position and angle of Camera R in the frame {**C**}, were estimated as follows. First, the pipette tip was moved randomly. Then, 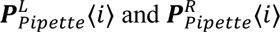 were obtained for the *i*-th movement. On the basis of these pairs of coordinates, the 3 × 3 essential matrix ***E*** that satisfies the constraint below was estimated using the “estimateEssentialMatrix” function,

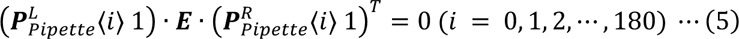

Then, the “relativeCameraPose” function was used to calculate the rotation and translation relative to Camera L from the essential matrix.

### 2. Transformation of the 3D pipette tip coordinates to the frame centered at the pivot point

In previous research that proposed a calibration method for the robot arm^31^, the edge of the arm was called the effector, and the coordinate frame centered at the effector was defined as frame {**E**}. Although the Hexapod six-axis stage was used instead of a robotic arm in the current study, the pivot point, which is the rotation center in the Hexapod control system, can be considered as the origin of frame {**E**} when it is fixed at a certain coordinate (Figure 3C). Therefore, the pivot point was regarded as the effector in the current study. The frame of the robot base system is represented as frame {**B**} (Figure 3C).

The goal of this section is to estimate 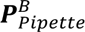, the coordinates of the pipette tip in the frame {**B**} (3 × 1 vector). 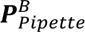 is expressed in two different ways. The first one is:

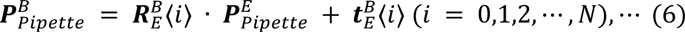

where 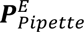, a 3 x 1 vector, is the initiation position of the pipette tip in frame {**E**}, and 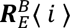 and 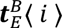 are the rotation and position of the effector in frame {**B**} after the *i*-th movement. 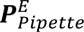 is the only unknown variable required to calculate 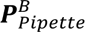 in equation (6). The second expression of 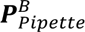 is:

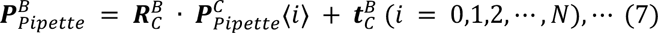

where 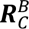 and 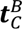 are the rotation matrix and translation vector in the camera coordinate system observed from frame {**B**}, respectively, and which are unknown here, and *N* is the total number of movements. 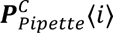 is the position of the pipette tip in frame {**C**} after the *i*-th movement. By solving Equations (6) and (7) simultaneously, the following equation is obtained:

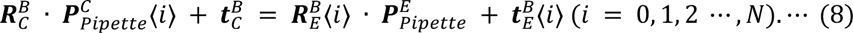

Although it is impossible to obtain 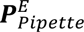 directly, the following equation (9) can be obtained by subtracting equation (8) at *i* = 0 from equation (8) at the *i*-th movement while keeping a fixed rotation of the end effector 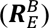, since 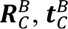 and 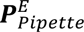 are constant:

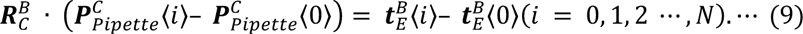

In equations (6–9), *N* was set at 200. Here, we can obtain 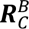 using singular value decomposition (SVD). Next, by subtracting equation (8) at *i* = 0 from equation (8) after the *i*-th movement with rotation, the following equation is obtained:

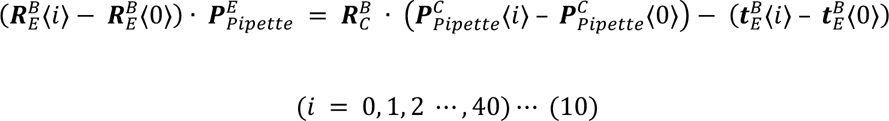

Then, 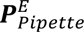 can be calculated using SVD. Finally, 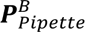 is estimated from equation (6).

### 3. Compensation for translation error

There are some error-inducing factors that were not discussed in the previous research. In the current study, two additional compensations for these errors were performed. The first error is the error derived from the pipette detection on the captured images and camera calibration, which mainly

affects the estimation of 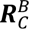 in equation (9). To better estimate 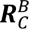 from error-laden point data in the stereo images, the robust estimation technique of MSAC was used. This algorithm consisted of five steps as follows:

Step 1: Determination of the sample number for estimation of 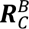. Since there are nine unknown variables in the 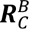 matrix, at least nine equations must be solved simultaneously in the estimation. Each pair consisting of the effector translation vector 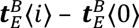 in frame {**B**} and the pipette translation vector 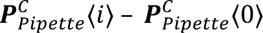 in frame {**C**} contributes three equations, one for each axis. Therefore, at least three pairs of translation vectors are necessary to estimate 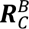. Considering the stability of the estimation, nine samples were randomly chosen from 200 samples.

Step 2: Estimate 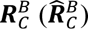 using the samples obtained in Step 1.

Step 3: Estimate the translation vector 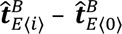 from 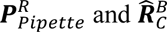.

Step 4: Estimate the cost function:

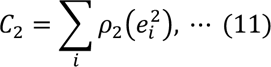

where

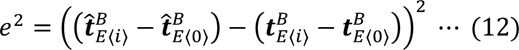

and *ρ*_1_ is defined as follows:

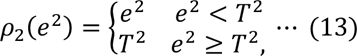

where *T* = 0.015 is a threshold for the errors. The samples above this value are called outliers and are virtually omitted from the estimation by replacing *e*^1^ with *T*^1^, which means ignoring outlying deviating points.

Step 5: Repeat Steps 1–4 and adopt 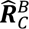, which minimizes the cost function.

From this procedure, the movement of the pipette tip was estimated from the captured images with an error rate of less than 15 μm for a 0.5 mm movement (Figure 3E).

### 4. Compensation for rotation error

The second aspect of the error is deformation of the hexapod robot due to its weight. During the system development, we found that when the pipette rotated around the pivot point that should correspond to the pipette tip, the tip location shifted a maximum of 0.4 mm (Figure 3G). The estimation of 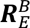 was not affected by this shift because the repeatability of the translation was stable as long as the angle of the stage was constant, but the estimation of 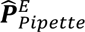 was biased and shifts of a few hundred micrometers were unacceptable for our purpose. This kind of error is commonly encountered when a commercial device is used. We handled this problem by grid searching of the pipette tip around the estimated 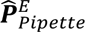 and measuring the shift of the pipette tip directly.

The shift of the pipette tip caused by the rotation was found to be invariant within the possible range of the injection position. Thus, the translation of the robot stage can be expressed as follows:

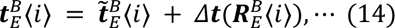

where 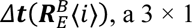 vector, represents the 3D shift between the expected position from the rotation 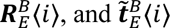 is the expected movement after the *i*-th robot movement. Without rotation,

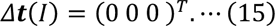

Considering these effects, equation (10) should be impossible to solve because there are two additional unknown terms, 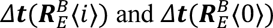:

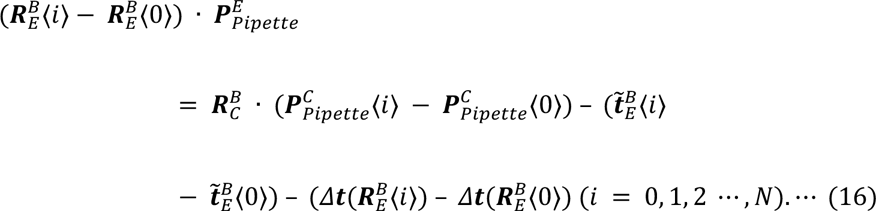

Using a single effector position can introduce bias into our estimate of 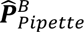. To address this, we conducted multiple estimations using 10 different effector positions. This approach allowed us to better infer the likely distribution of 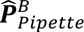, which we found to vary within a cube with sides measuring approximately 0.7–0.8 mm (Figure S4E). To align the pipette tip more precisely with the pivot point, the effector was positioned within a 0.8 mm square cube. The aim of this section is to find the position that minimizes the shift of the pipette tip position during rotation. The procedure consists of eight steps as follows:

Step 1: Start by creating a grid of 64 evenly spaced points within a cube of side *x* mm. Initially, *x* = 0.8 mm (Figure S4F).

Step 2: Move the effector to the *i*-th point within the grid.

Step 3: Rotate the stage by –7° around the Y axis (referred to as the V rotation), and by –15° around the Z axis (referred to as the W rotation).

Step 4: Measure the pipette shift for both the V and W rotations, denoted as *e*_*v*_ and *e*_*w*_, respectively.

Step 5: Calculate the squared error for the *i*-th position using the formula 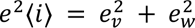.

Step 6: Repeat Steps 2–5 for all 64 points in the grid and identify the position where *e*^1^〈*i*〉 is minimized.

Step 7: Narrow the search grid to a cube centered around the point identified in Step 6. The new *x* is set to two-thirds of the previous *x* (Figure S4G). Return to Step 1 and repeat the process.

Step 8: Repeat Steps 1–7 over four cycles. This allows the possible volume where the pipette tip could exist to be narrowed to a 0.16 mm square cube (Figure S4H).

Performing these procedures decreased the shift to less than 0.03 mm at the rotation that was used for the calibration. However, the error at the maximum of 0.3 mm was still produced at the other angles (Figure 3G). Finally, we measured 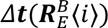 in a grid-based approach for both the V and W rotations, and calculated regression models to estimate 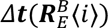 from V and W rotations. First, 40 × 40 pairs of V and W grid search points were arranged within the ranges of ± 14° for V rotation and ± 23° for W rotation. The number of actual sets of angles was approximately 1000, excluding those that could not be moved due to the physical constraints of the robot. The pipette tip movement was measured twice at each search point and averaged. Then, we used the thin plate spline method to construct a regression model,

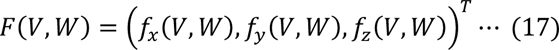

where *f*_*x*_(*V*, *W*), *f*_*y*_(*V*, *W*), and *f*_*z*_(*V*, *W*) represent the functions of (*V*, *W*) to output the shift by rotation of *V* and *W* angles in the *X*, *Y*, and *Z* axes, respectively (Figure 3H). The shift that was estimated from this model was compensated for by translation of −*F*(*V*, *W*). Thus, the final error can be written as follows:

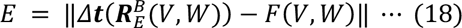

By developing these procedures, 99.8% of the shift caused by the rotation was less than 50 μm and the average shift was 0.021 ± 0.0002 mm (n = 1274) at any position and any angle within the range of the injection experiments (Figure 3J, K and Figure S4I, J).

### Registration of laser reflection point to the pipette tip and laser scanning on the cortical surface

A 26G needle (NN-2613S; Terumo Corporation, Tokyo, Japan) was cut to make a hollow metal rod that was used as the reference point to align the laser reflection point with the pipette tip in the robot base system. The cut needle was fixed in a position slightly above the base plate. The glass pipette tip was moved to the center of the metal rod with the robot, and the robot coordinates at this point were recorded. Next, after the pipette was moved up by the actuator, the laser point was matched to the center of the metal rod by translating the robot stage position, and the coordinates of this point in the robot base system were recorded. From these two coordinates, the transformation matrix of the laser reflection point and pipette tip was obtained.

The laser reflection point was moved to the bregma by eye inspection using Camera S. An area of ± 5 mm × ± 4.5 mm (anterior-posterior [AP] axis × medial-lateral [ML] axis) centered on the bregma was scanned, with surface points measured every 0.2 mm along the AP axis and every 0.13 mm along the ML axis. The AP axis corresponded to the Z axis in frame {**B**} and the ML axis corresponded to the Y axis in frame {**B**}. In the mouse brain, the Allen Mouse Brain Atlas CCFv3 3D template data (https://connectivity.brain-map.org/static/referencedata) were fitted to the laser-scanned 3D data using an affine transformation including rotation, translation, and anisotropic scaling. To minimize the difference between the coordinate along the DV axis (X axis in frame {**B**}) measured by laser scanning and the X coordinate predicted from the transformed template data, the affine transformation parameters were repeatedly updated. The 3D interpolation model based on the 3D template data after the affine transformation was calculated using the “scatteredInterpolant” function.

In the marmoset brain, nine parameters of the quadric surface, *P_ij_* (*i* = 0, 1, 2; *j* = 0, 1, 2), were defined by the following equation:

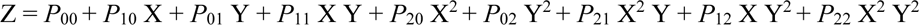

These parameters were estimated from the laser-measured data using the robust least squares method. The difference between the X coordinate measured by laser scanning and the X coordinate predicted from the estimated surface was calculated, and the top 1% and bottom 1% of the distribution were removed as outliers. A 3D interpolation model of the remaining measurement data was constructed using the “scatteredInterpolant” function.

### Acquisition of cortical surface images by Camera S

The 2D coordinates of the pipette tip near the center of the image captured by Camera S were recorded. When the cortical surface was imaged with Camera S, the pipette was moved up with the actuator. In the mouse brain, the image was acquired at 0.6 mm intervals 64 times and a total area of approximately 8 × 8 mm was covered (Figure S1A). The X coordinate on the cortical surface with the Y and Z coordinates of one of the predetermined 64 YZ points was calculated from the 3D interpolation model obtained in the previous step (Figure S1A) and the robot was moved so that the point that corresponded to the pipette tip coincided with the position to acquire the image (red point in Figure S1B). In the marmoset, the image resolution was two-thirds lower than that in the mouse brain; images were acquired at 1 mm intervals 106 times, and a total area of approximately 9.8 × 17.3 mm was covered.

### Building the neural network for vessel segmentation

For each image, the area of 3.8 × 3.8 mm centered at the pipette tip coordinates was cropped and the cropped image was used as the input image for SA-UNet. For training of SA-UNet, twenty images of retinal fundus vasculature from the Digital Retinal Images for Vessel Extraction (DRIVE) dataset were initially prepared, and six original images and labels of cortical vessels in mouse brains were further prepared. This model worked well for relatively narrow vessels but not for bold vessels or poorly conditioned surfaces. Several images were therefore added to the training dataset. The methodology for choosing more “informative” images to add to the training dataset is called active learning. EquAL^27^, which calculates the difference in prediction between the original and inverted images as “consistency” of the inference, was adopted as a criterion. In addition, the training annotations were based on the prediction of the neural network with manual modifications in a process called CNN-corrected segmentation^28^. This EquAL selection and CNN-corrected segmentation cycle was repeated three times with 33 original images (six images were manually annotated and 27 images were CNN-corrected) and 20 DRIVE dataset images being used to train the neural network. The final metrics of the modified SA-UNet were as follows: MCC, 0.713; F1, 0.779; AUC, 0.951; specificity, 0.874; sensitivity, 0.907; and accuracy, 0.881. All training processes were written and performed in Python and Keras.

### Transformation of the original stitched image to the YZ stitched image

First, 64 vessel segmentation images were stitched with the automatic panoramic image stitching method (Autostitch.exe^53^). Thus, in each mouse, one large area of approximately 8 × 8 mm was obtained as the original stitched image. To map the target injection sites determined in the 2D stitched image to the 3D curvature of the cortical surface, it is necessary to compensate for image distortion caused by the difference between the camera angle and laser emission angle. The 3D shape of the cortical surface estimated from the laser scanning was rotated through a combination of V and W rotation parameters (*v*, *w*). The coordinates of the pipette tip (reference point) at each image acquisition, which were assumed to be on the cortical surface, were also rotated by (*v*, *w*), and these points were projected to the 2D plane along the corresponding rotation axis. The geometric transformation from the projected 2D coordinates of the pipette tip to the pipette tip coordinates in the original stitched image was performed using the “fitgeotrans” function. Then, the geometric transformation, i.e., the distance between the transformed point of the pipette tip point and the corresponding point in the stitched image, was calculated. This distance was summed over all acquired images to represent the error. Then, after changing the combination of rotation parameters, the error was calculated again. These procedures were repeated to find the combination of rotation parameters that minimized the error. The combination of the rotation parameters that minimized the error was (–17.47 ± 0.19°, 5.09 ± 0.51°) in seven mice and (–18.18°, 3.241°) in one marmoset.

Using the rotation and its geometric transformation that minimized the error, the vessel structures in the original stitched image were mapped onto the 3D curvature of the cortical surface. Finally, this 3D image was projected into the YZ plane (YZ stitched image) to determine the multiple injection sites.

### Injection sites planning algorithm

If the virus was injected into the centers of the hexagons of the 2D honeycomb structure as the 2D closest packed structure, the gene was assumed to be expressed at the highest filling rate (or uniformly expressed in space). If the distance between the centers is *d*, the area of the hexagon with a side of 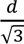 is expressed 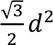 (desired single-injection area). If the distance from each injection site to its nearest injection site was *d*, and the total area of the cortical region including the safety region and the other regions (blood vessel and safety margin regions) was *A*, the number of injection sites *N*_t_ was calculated as approximately 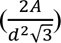. However, the injection sites could not be set in blood vessel or safety margin regions. Therefore, we determined the distribution of injection sites by repeating the clustering method to approach a honeycomb structure. First, the initial number of sites was set to *N_t_* + 10. Then, the safety region was divided into (*N_t_* + 10) clusters using k-means clustering and the centroid of each cluster was determined as the injection site candidate. Each

cortical pixel within the region including the safety region and the other regions was assigned to the

nearest candidate. Then, the total cortical area assigned to each final injection site candidate was calculated as the assigned injection area. If the median value of the assigned injection area did not exceed the desired injection area, the two sites with the smallest and second smallest assigned injection areas were removed and a new candidate site was added. Then, the clustering was performed again. This procedure was repeated approximately 10 times until the median value of the assigned injection area exceeded the desired injection area. Based on the misalignment distribution and the simulation of the bleeding risk probability, the safety margin distance was generally set at 65 µm. *d* was set to 0.35–0.65 mm.

### Sequential injections into the target injection sites

The order of multiple injection sites was determined by solving the traveling salesman problem. The order of the target points near the bregma was set to be earlier than the order of the points around the edge of the craniotomy. The series of pipette tip movements were made according to the following steps:

Step 1: Stay 3 mm above the bregma.

Step 2: Move to 1 mm above the target injection point on the cortical surface. Step 3: Move 0.2 mm above the target point at a speed of 20 mm/s.

Step 4: Change the speed to 5 mm/s.

Step 5: Change the angle of the pipette. At this point, a new coordinate system in frame {**E**} was set. The X axis of the device was aligned with the vertical line to the cortical surface, and the position was moved to a point 2 mm vertically above the target point.

Step 6: In the mouse, the pipette was transiently inserted to a depth of 0.6–0.8 mm at a speed of 6 mm/s, penetrating the dura mater. Then, the pipette was pulled to the target depth (0.3 mm) at 0.1 mm/s and held for a time setting of 15–30 s to inject the solution. In the marmoset, the pipette was inserted to the target depth (0.5 mm) at a speed of 0.1 mm/s and held for a time setting of 5–10 s to inject the solution.

Step 7: Move to 0.2 mm vertically above the insertion point (at the speed of 0.1 mm/s). Step 8: Move to 1 mm vertically above the injection point (20 mm/s).

Step 9: Move to 4 mm above the injection point and reset the angle.

The cycle of Steps 2–9 was repeated for subsequent injection sites.

### Animals

All animal experiments were approved by the Animal Experimental Committee of the University of Tokyo. C57BL/6 mice (males and females, 8–14 weeks old) were used. All mice were provided with food and water ad libitum, and housed in a 12:12 hour light-dark cycle, unless otherwise mentioned. One laboratory-bred common marmoset (Callithrix jacchus, female, 2 years and 5 months old) was also used in the present study. The marmoset was kept on a 12:12 hour light-dark cycle and was not used for other experiments prior to the present study.

### Head plate implantation for mice

Head plate implantation was performed as the protocol described previously^37^. Mice were anesthetized by intraperitoneal injection of a mixture of ketamine (74 mg/kg) and xylazine (10 mg/kg), followed by topical administration of eye ointment (Tarivid; 0.3% w/v ofloxacin, Santen Pharmaceutical, Japan) to prevent eye-drying and infection. During surgery, body temperature was maintained at 36°C–37°C with a heating pad. The head of each mouse was sterilized with 70% ethanol, the hair was shaved, and the skin covering the neocortex was incised. The exposed skull was cleaned, and a head plate (Tsukasa Giken, Japan) was attached to the skull using dental cement (Fuji lute BC; GC, Tokyo, Japan; and Bistite II; Tokuyama Dental, Japan). The surface of the intact skull was coated with dental adhesive resin cement (Super bond; Sun Medical, Japan) to prevent drying. An isotonic saline solution containing 5 w/v% glucose and the anti-inflammatory analgesic carprofen (5 mg/kg, Remadile; Zoetis, NJ) was injected intraperitoneally after all surgical procedures. The number of mice per cage was 2–5 before the head plate was attached, and then after attachment the mice were housed singly to avoid damage to the head plate and glass window.

### Head plate implantation in the marmoset

All surgical procedures on the marmoset were performed under aseptic conditions, as described previously^15^. Following anesthetization, the marmoset was placed in a stereotaxic instrument (SR-6C-HT; Narishige, Japan), with anesthesia maintained by inhalation of isoflurane (1.5–4.0% in oxygen). Oxygen saturation (SpO2), heart rate, and rectal temperature were monitored continuously. The antibiotic ampicillin (16.7 mg/kg) and the antiemetic maropitant (1000 mg/kg) were administered intramuscularly. The anti-inflammatory drug carprofen (4.4 mg/kg) was intramuscularly injected to reduce pain and inflammation in the perioperative period. Acetated Ringer’s solution (10 mL) containing riboflavin sodium phosphate (200 µg) was also administered subcutaneously. Hair was removed from the head using a depilatory, and the head was sterilized with povidone iodine, followed by exposure of the skull. Lidocaine jelly was applied to wound sites to reduce pain. A head plate was attached to the skull using universal primer (Tokuyama Dental, Japan), dual-cured adhesive resin cement (Estecem II; Tokuyama Dental), and dental resin cement (Superbond; Sun Medical, Japan).

### Virus production

pGP-AAV-syn-jGCaMP7c variant 1513-WPRE was a gift from Douglas Kim & GENIE Project (Addgene viral prep # 105321-AAV1; http://n2t.net/addgene:105321; RRID: Addgene_105321)^54^. The AAV plasmid of human synapsin I promoter (hSyn)-tetracycline-controlled transactivator 2 (tTA2) was constructed by subcloning the DNA fragments containing hSyn and tTA2 into pAAV-MCS (Agilent Technologies, CA, USA). The AAV vector was produced as described previously^15,35,55^. For the generation of pAAV-TRE-GcaMP6s-P2A-GcaMP6s-WPRE (tandem GcaMP6s), gene fragments of GcaMP6s and P2A were obtained from pGP-CMV-GcaMP6s-WPRE (which was a gift from Douglas Kim & GENIE Project [Addgene plasmid # 40753; http://n2t.net/addgene:40753; RRID: Addgene_40753]), and pAAV-hSyn1-GcaMP6s-P2A-nls-dTomato (which was a gift from Jonathan Ting [Addgene plasmid # 51084; http://n2t.net/addgene:51084; RRID: Addgene_51084]), respectively^8,34^. pAAV-TRE-GcaMP6f-

WPRE was used as a template for this plasmid. AAV plasmids were packaged into AAV serotype 9 using the AAV Helper-Free system (Agilent Technologies). In brief, pAAV vector, pRC9, and pHelper plasmids were transfected into HEK293 cells. Seventy-two hours after transfection, AAV2/9 particles were purified using the AAV Purification kit (Takara, Japan). The AAV solution was concentrated to the optimal volume by centrifugation using an Amicon Ultra-4 100k centrifugal filter unit (Millipore). The number of genomic copies was quantified with intercalating dyes (Thermo Fisher Scientific, MA, USA) and two sets of primers for WPRE or hGHpA genes, using LightCycler 480 (Roche, Basel, Switzerland). The final titration of the AAV was estimated as relative quantitation according to a calibration curve calculated from the known numbers of copies of AAV plasmids.

### Virus injections into the mouse cerebral cortex

Following anesthetization, a craniotomy measuring 8 × 8 mm and centered at the bregma was made in the skull of each mouse, with the dura mater being left as intact as possible. Anesthesia was maintained throughout surgery by inhalation of 1% isoflurane. To prevent cerebral edema, dexamethasone sodium phosphate (1.32 mg/kg; Decadron, Aspen Japan, Japan) was administered intraperitoneally 30 minutes before the start of the craniotomy. The AAV solution (AAV1-syn-jGCaMP7c; 7.86 × 10^12^ vg/mL) was subsequently injected using a pulled quartz-glass capillary (broken and beveled to an outer diameter of 25–30 µm; Nakahara Opto-Electronics Laboratories, Inc., Japan) and an injector (Nanoject III). The glass capillary was filled with mineral oil (Nacalai Tesque, Japan) and the solution of virus was drawn up from the tip prior to injection. To validate the accuracy of the robotic manipulation, fluorescent dye (Vybrant^TM^ CM-DiI Cell-Labeling Solution; Invitrogen, CA, USA) was injected instead of virus solution. The pipette was inserted into the target (cortical depth of 500 µm) and held for 15 s, and 3 nL of the dye stock solution were injected into each site at a rate of 1 nL/s with a 1 s interval, followed by a 30 s wait

until the next injection. After all injections were finished, a mixture of ketamine (74 mg/kg) and

xylazine (10 mg/kg) was administered intraperitoneally, and the cortical surface was viewed with an epi-fluorescent microscope (THT macroscope; Brainvision, Japan) to determine the fluorescent spots. Subsequently, the animals were deeply anesthetized for slice preparation.

### Virus injections into the frontoparietal cortex in the marmoset

For wide-field one-photon calcium imaging of a marmoset, a rectangular craniotomy of 8 × 16 mm was made over the right dorsal cortex from area 8A to area PE, and the dura mater was removed. The AAV in the pipette was injected as described above. The virus solution consisted of AAV9-hsyn-tTA (5.38 × 10^12^ vg/mL) and AAV9-TRE3G-tandem-GcaMP6s (1.365 × 10^13^ vg/mL). The pipette was inserted to a depth of 500 µm from the cortical surface and held for 2.5–5 s, and the virus solution was injected. A total of 30 nL of solution was injected into each site at a rate of 2 nL/s with a 5 s delay after each injection. A rectangular glass coverslip of 15 × 8 mm (approximately 150 µm thick; Matsunami Glass, Japan) that was attached to the bottom of a polyacetal chamber (height, 1.2 mm) with UV-curing optical adhesive (NOR-61) was pressed onto the cortical surface, and the space between the chamber and the cortical surface was filled with dental cement (Fuji Lute BC; GC, Japan). The edge of the chamber was sealed with dental adhesive resin cement (Super bond; Sun Medical, Japan). An area of 14.2 × 7.2 mm, excluding the width of the chamber’s frame, formed the field of view for cortical imaging. The marmoset was allowed to recover for 7 weeks prior to imaging.

### Measurement of the insertion points and depths in the mouse cortex

The location of each insertion was observed as CM-DiI fluorescence using an epifluorescence microscope. Elastic and consistent image registration using bUnwarpJ^56^ was applied to the fluorescence and surface vascular pattern images to determine the injection sites. Because the registration was not accurate at the periphery of the images, the images were cropped around the targeted injection site before application of the algorithm. The distance between the center of fluorescence and the planned injection site was measured on the registered image using Fiji^57^. If the insertion point was observed as a shadow surrounded by fluorescence, the center of the observed shadow was used for measurement, instead of the center of fluorescence.

The trajectory of the pipette insertion within the cortex was traced according to the fluorescence derived from CM-DiI in the slice. Each mouse was deeply anesthetized with a mixture of ketamine and xylazine, and then transcardially perfused with PBS (10 mL) followed by 4% formaldehyde (30 mL). Their brains were post-fixed with the same fixative overnight at 4°C and then coronally sectioned at a width of 100 µm. The sections were counterstained with a fluorescent Nissl stain (NeuroTrace blue; 1:500; Invitrogen). Images were acquired with an epifluorescence microscope (BX53F; Olympus, Tokyo, Japan) and a digital CCD camera (RETIGA2000R; Qimaging, Canada). Since the plane of the slice and the trajectory of the injection did not always coincide, fluorescence derived from the same injection was sometimes observed across multiple slices. Therefore, the putative depth of the pipette tip was estimated as follows. First, the distance from the deepest point of the pipette track (observed as fluorescence) to the surface was measured for each slice. If the pipette trajectory was visible as a shadow surrounded by fluorescence, the distance from the deepest point of the shadow to the surface was measured. Adjacent slices were compared to determine whether the trajectories originated from the same injection. These processes were performed using Fiji^57^.

### Estimation of bleeding risk probability

The distribution of misalignment of the actual injection sites from the target injection sites (82 sites from three mice) was fitted using a non-parametric Gaussian kernel restricted to positive values. Based on the data, the following methods were used to determine the relationships between the distance of the injection sites from blood vessels and the hemorrhage probability. First, in one cerebral cortex image obtained with Camera S, the distance between each target injection site was set to 0.4 mm and the total number of injection sites per trial was set to approximately 120. The safety margin distance (the minimum distance of the target site from blood vessels), *D*, was specified (0–80 μm). In this condition, using the clustering methods as described, the target injection sites were determined. Next, to determine the simulated injection site, the position of the target site was shifted according to the estimated misalignment distribution. Those simulated injection sites that hit a vessel were considered as bleeding sites and their number was counted. The simulation was conducted 100 times for each *D* value. Finally, the proportion of the simulated injection sites that hit the vessels was averaged for each *D* value.

### Comparison of the blood vessels using microscopy and vessel segmentation

To examine the limits of the ability of the modified SA U-Net to extract blood vessels from the camera image, the results of the vessel segmentation were compared with vessels observed under 5× epi-illumination microscopy. One microscopic image of 1 × 1 mm size was taken and vessels were masked manually. The following process written in MATLAB (R2021b) was then used to measure the thickness. 1. Distance transformation was applied to the vessel mask. The brightness values of the pixels in the mask were determined according to their distance from the nearest non-zero pixels. 2. The vessel mask was skeletonized to obtain the furthest line from the boundary. 3. The distance map and the skeletonized vessels were multiplied. This yielded a skeleton of the blood vessel mask where the pixel brightness corresponded to the vessel thickness (skeletonized distance map). The vessel mask obtained by the modified SA U-Net was also prepared for this analysis. The actual output from the modified SA U-Net is an 8-bit grayscale image, and this was binarized by applying the threshold (205) that was used to determine the injection site. Then, projective transformation was applied to the segmentation result to match the microscopic image using manually extracted feature points. The thickness that the modified SA U-Net was able to recognize was obtained by calculating the overlaid area between the segmentation and the skeletonized distance map.

### Wide-field calcium imaging of the marmoset cortex

In the marmoset, images were acquired using a fluorescent zoom macroscope equipped with a high-numerical aperture macro lens (PlanNEOFLUAR Z 1.0×, NA 0.25; Carl-Zeiss, Germany) and an EM-CCD camera (iXon Ultra 888; Andor Technology, UK). The microscope body with the objective was rotated 16.5° around the anterior-posterior axis and the marmoset chair was tilted 7° so that the optical axis was nearly perpendicular to the glass window. Images of 512 × 512 pixels were captured using a Solis camera system (Andor Technology) at a sampling rate of 30 frames/s. Optical zoom magnification was set to cover the entire window in the marmoset. The center wavelength of the LED for the wide-field calcium imaging was set to 470 nm, and the light intensity under the device was 6 mW. The animal was trained to sit in an awake head-fixed condition, and the position of a spout was adjusted so that the marmoset could lick it. A series of 18 000 images were acquired while 40 µL of sucrose water was delivered as a reward every 10–20 s. No apparent fading of the fluorescent signal was observed over a duration of several days under this experimental condition.

### Two-photon calcium imaging of the marmoset cortex

The marmoset cortex was imaged with a custom-built two-photon microscopy (Olympus) system equipped with a water immersion objective lens (Olympus XLPLN10XSVMP, NA 0.6; working distance 8 mm) and an Nd-based fiber-delivered femtosecond laser (Femtolite FD/J-FD-500, pulse width of 191–194 fs, repetition rate of 51 MHz; IMRA, USA) at a wavelength of 920 nm. Details of this implementation were described previously^15^. The optical axis was adjusted to be nearly perpendicular to the cranial window by tilting the microscope body 4.8° and the marmoset chair 5°. To shield the microscope objective from possible stray light, an aluminum foil dish was attached to the implanted metal chamber using silicone elastomer (Dent Silicone-V, Japan), and the space over the animal’s head was covered with lightproof cloth. The head-fixed animal was habituated under the microscope. A series of 18 000 images were acquired while 40 µL of sucrose water was delivered as a reward every 10–20 s. The imaging field area was 636 × 636 µm and 512 × 512 pixels. Two different regions were imaged on different days.

### Identification of cortical areas in the marmoset

To infer the imaging areas of the marmoset frontoparietal cortex, intracortical microstimulation was performed 1 week after the imaging experiments, as described previously^35^. The glass window was removed and a tungsten microelectrode with an impedance of 0.5 MΩ (World Precision Instruments, Sarasota, FL, USA) was inserted to a depth of 1.5 mm from the cortical surface, and a train of 15 cathodal pulses (0.2 ms duration at 333 Hz) were applied. Current ranging from 10 to 100 μA was sequentially applied for each region. The minimum current required to elicit body part movements, as confirmed by an expert experimenter, was defined as the threshold current for that stimulation site. The type of body part movement (extension, flexion, or rotation) induced was determined by visual inspection by the experimenters. The cortical map in Figure 5D and Figure S6A was determined as follows. First, we identified the cortical area that had a relatively low movement threshold in comparison with more posterior or anterior areas. We considered this area to be the main part of M1. Then, the border between M1 and the caudal part of the dorsal premotor cortex (area 6DC) in this animal was defined as the line at which the threshold current apparently increased along the posterior-to-anterior direction. The marmoset cortical map by Paxinos^58^ was aligned with the dorsal cortex of this animal so that the midline and the border between M1 and area 6DC matched between the Paxinos map and this animal.

### Monitoring licking and body movement

Two CMOS cameras (DMK33UP1300; ImagingSource, Taipei, Taiwan) equipped with a fixed-focus lens (focal length, 3.0 mm; #89-341; Edmund, NJ, USA) and a varifocal lens (focal length, 2.7–12.0 mm; Spacecom, Japan) were set at a distance of 270 mm and an angle of 35° from the front of the marmoset face. During the imaging experiment, two 480 × 480-pixel images were simultaneously obtained at 50 frames/s. Licking was detected as a change in the average luminance within a region of interest (ROI) set around the mouth using ImageJ software (National Institute of Health, USA), followed by processing in MATLAB (R2022a). A high-pass filter was applied to the signal to remove baseline trends and it was dichotomized at a threshold of two standard deviations.

### Processing of wide-field calcium imaging data for the marmoset

The data obtained from the marmoset across three sessions on different days were analyzed using custom MATLAB routines (R2021a, R2022a; MathWorks, USA). Fluorescence data were obtained as 512 × 512-pixel tiff images, which were cropped to 241 × 442-pixel images prior to further processing. The time-varying baseline fluorescence of each pixel (*F*) was estimated for a given time point as the 10^th^ percentile value over 30 s around it, and the relative fluorescence change (Δ*F*/*F*) of each pixel was calculated. Because three sessions were performed on different days, the images were registered across sessions. The average frames of each imaging session were used for registration. The averaged frame of the first session was regarded as the reference image for alignment. A geometric transformation matrix was generated to align the average frames of the second and third sessions to the reference image.

To assess the functional connectivity that was not directly related to movements, we extracted the neuronal activity during the quiet state, which we defined as the time period excluding that from 0.3 s before to 2 s after the occurrence of licking or body movement. Then, correlations in the fluorescence change between the areas during this quiet state were calculated. Hierarchical clustering was applied to the resulting correlation matrix, using the Calinski-Harabasz criterion, and two distinct clusters were identified. Seed correlation maps were created using the ROIs of areas PFG and 4ab as seed points. The correlation coefficients between these seeds and all other pixels were visualized as seed correlation maps.

A linear regression model was constructed using a design matrix composed of multiple variables. Each variable was generated as follows. For the “body_analog” measurement, the absolute difference between two adjacent frames was calculated, and its average within that frame was regarded as the motion intensity at that time. The z-scored value of the motion intensity was assigned to the “body_analog” variable. The “body” variable was binarized “body_analog” data with a threshold value of 2. For the “eye_analog” measurement, ROIs were placed around the pupils of both eyes, and movement traces were extracted using DeepLabCut^59^. The position of each eye was then determined by calculating the average position of the ROIs around its pupil. To quantify the intensity of eye motion, the speed of each eye’s movements was averaged. This calculated intensity was then normalized using z-scoring and recorded as the “eye_analog” variable. The “eye” variable was derived from this “eye_analog” data, binarized with a threshold value of two. For the “eyelid_analog” measurement, ROIs were placed on eyelids of both eyes, and movement traces were extracted using DeepLabCut. Subsequently, the z-scored values of their vertical coordinates were averaged and assigned to the “eyelid_analog” variable. For the “jaw_analog” measurement, an ROI was placed on the jaw and the movement traces of the jaw were extracted using DeepLabCut. Then, the speed was calculated to quantify the intensity of jaw motion. The z-scored value of this intensity was assigned to the “jaw_analog” variable. The “jaw” variable was derived from the “jaw_analog” data, binarized with a threshold value of 0. For the “lick” measurement, an ROI was placed on the tongue, and its appearances were extracted using DeepLabCut. The times when the tongue was recognized were assigned a value of 1, while all other times were assigned a value of 0. For each vector of the “body”, “eye”, “jaw”, and “lick” variables, time-shifted copies were generated, covering frames from 1 s before to 3 s after each event. For the reward variable, a vector containing a pulse at the time of reward delivery was copied in time, spanning the frames from the timing of the reward delivery to 3 s after each event. The “body_analog”, “eye_analog”, “eyelid_analog”, and “jaw_analog” variables were not subjected to time-shifting.

A ten-fold cross-validated linear regression was used to fit the design matrix to the fluorescence signals of each pixel, using ridge regularization and stochastic gradient descent solver. Subsequently, the variance explained (*R*^2^) by the full model was calculated. The selection of lambda for ridge regression was as follows: Fifteen candidate lambda values that ranged from 10^-4^ to 10^-1^ on a logarithmic scale were used to predict the time series data of a representative pixel’s fluorescence signal in a 10-fold ridge regression model. We then plotted the mean squared error and frequency of non-zero coefficients for each lambda value. The value of 0.0139 was chosen as it adequately balanced the model’s accuracy and the sparsity of coefficients. We confirmed the adequacy of this lambda value by verifying that the mean squared errors for other pixels were also comparably low. To estimate the explanatory power of each variable, shuffled models were created by randomizing all time points for only the specified variable group. The difference in the explained variance between the full and the shuffled model was considered as the unique contribution (Δ*R*^2^) of that variable group. The “mouth” variable group included the “jaw”, “jaw_analog”, and “lick” variables, while the “body”, “eye”, and “eyelid” groups included their respective analogue and binarized variables.

### Processing of two-photon calcium imaging data for the marmoset

The observed images were registered with rigid motion correction using NoRMCorre^60^ and their contrast was adjusted in field 1 and field 2. Then, the CNMF algorithm^61^ was applied to each set of data to obtain ROIs of active neuronal somata. The relative change in fluorescence for each ROI at a time point *t* was calculated 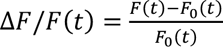, where *F*(*t*) was the mean of the fluorescence intensity value of the pixels within each ROI at *t*, and *F*_0_(*t*) was the baseline fluorescence at *t* calculated as the 8^th^ percentile of the *F*(*t*) value across *t* ± 15 s. The max Δ *F*/*F* values of the ROIs were statistically compared between two regions.

Linear regression models were constructed to analyze the relationships between the time-shifted neuronal activity and each behavioral variable (licking and body movement). To capture the potential influence of neuronal activity on the behavior, and its temporal dynamics, we created time-shifted versions of the activity data for each neuron. For each session, we used a time frame range from –30 to +30 (or ± 1 s range) to shift the activity data. This process resulted in 61 time-shifted traces for each neuron, including the original time point. These shifted traces were then aligned to form a new matrix for each neuron, with dimensions of 18 000 frames by 61 shifted activity traces. A linear regression model was fitted for each neuron, using its shifted activity matrix as predictors and the behavioral variable as the response variable. For licking decoding, a Gaussian filter with a 30-frame window was applied to the binarized lick signal and the filtered signal was used as the response variable. For body movement decoding, the motion intensity was used as the response variable. The predictive power of each model was assessed using *R*², which quantified the variance in the behavioral variable explained by the neuronal activity.

### Statistics

Statistical analyses were performed using MATLAB (R2021a, R2022a). Comparisons were made using Welch’s test, and one-way ANOVA followed by post-hoc one-sided Student’s *t*-test with Bonferroni correction. No statistical tests were run to predetermine the sample size. Data are presented as mean ± s.e.m., unless otherwise noted. Blinding and randomization were not performed.

## Data availability

The data supporting the findings of this study are available from the corresponding author upon reasonable request. All computer codes to realize the ARViS have been deposited at https://github.com/nomurshin/ARViS_Automated_Robotic_Virus_injection_System, and are publicly available.

