## Supplementary material for "ARViS: A bleed-free multi-site automated injection robot for accurate, fast, and dense delivery of virus to mouse and marmoset brains": Figures S1-6 and Table S1

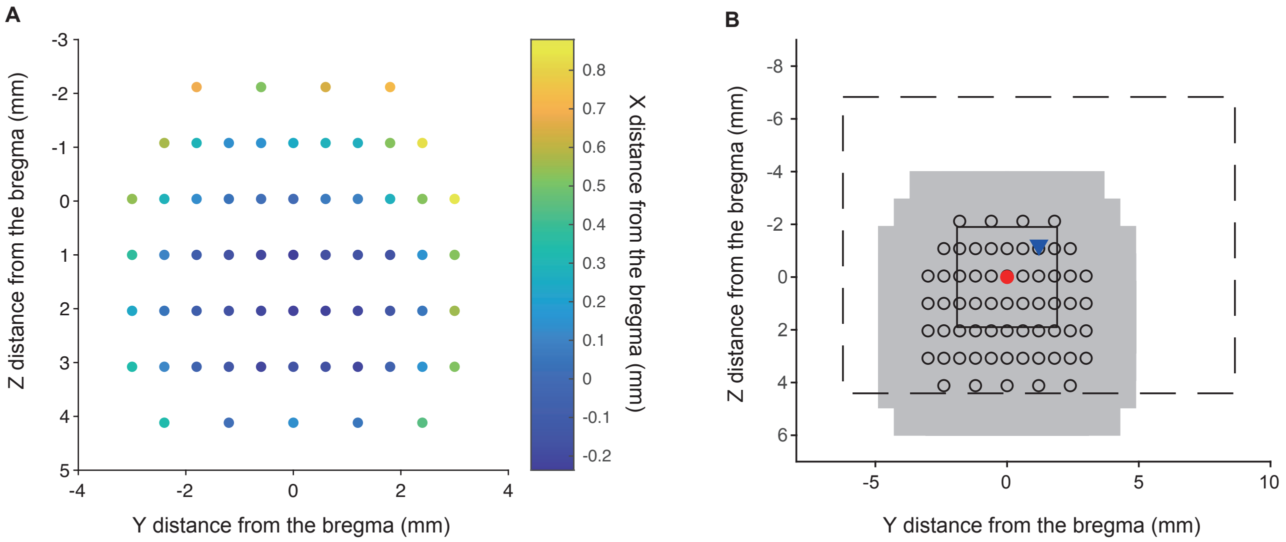


**Figure S1. Relationship between the image acquisition points and the photographic field of view.**

(A) Representative plots of the image acquisition points (dots) for mice. Sixty-two points were arranged on the surface measured by the laser distance sensor. The pseudo-colored bar shows the measured X coordinate of each point relative to the bregma.

(B) Schematic diagram illustrating the relationship between image acquisition points and the corresponding frames for SA-UNet inputs. Open circles denote all the image acquisition points shown in (A). For an image captured at the position indicated by the red filled circle, the resulting field of view from Camera S is delineated by the dashed frame. The blue triangle marks the center of this field of view. The red filled circle also signifies the pipette tip position within the corresponding field of view of Camera S when the pipette tip was moved down at the focal plane of Camera S. The solid black rectangle was the input image of this field of view for SA-UNet. The gray shading represents the area of the image where all 62 SA-UNet output images were stitched together (original stitched image).

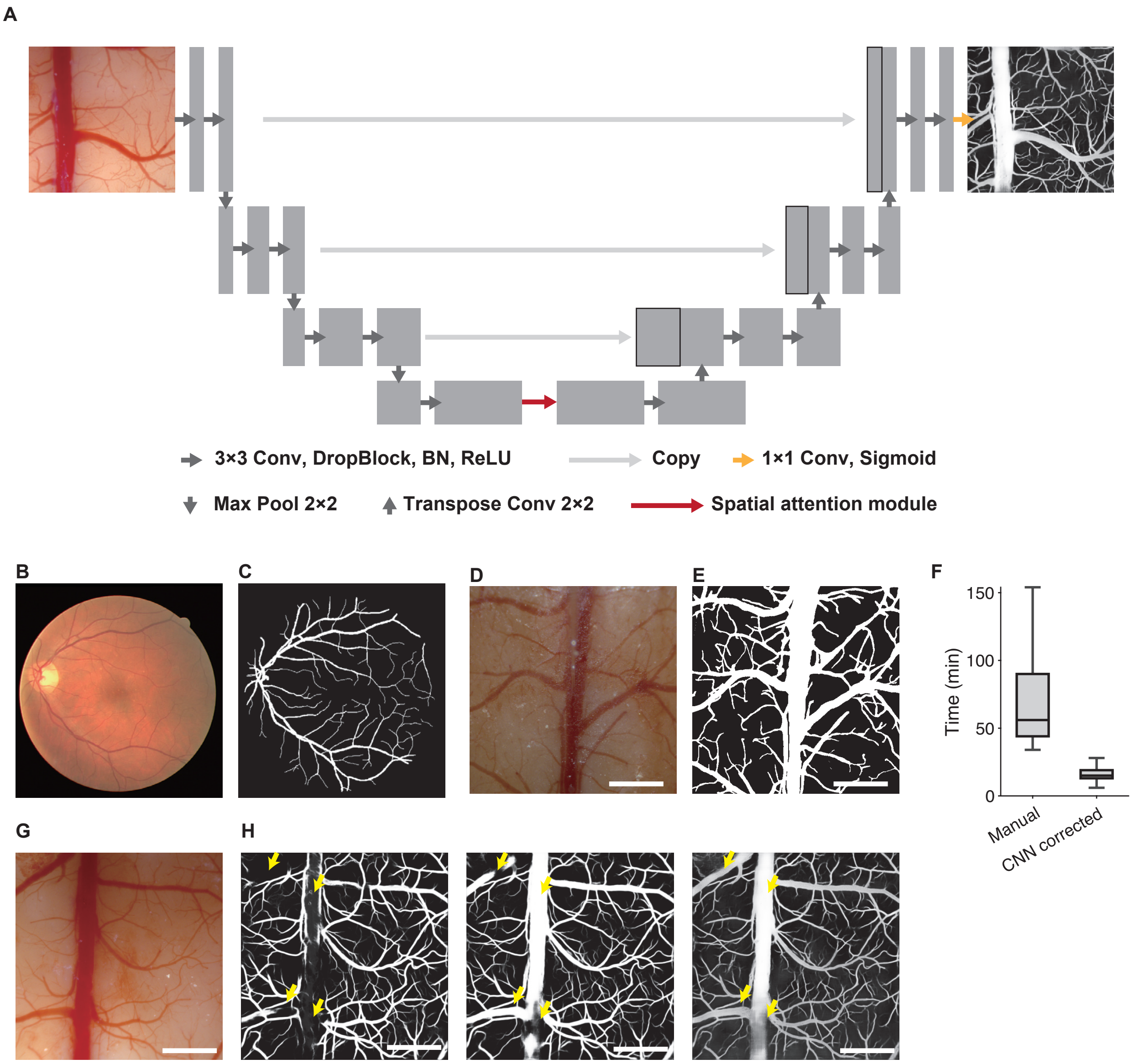


**Figure S2. Building the neural network for the vessel segmentation.**
(A) Schematic of the architecture of SA-UNet (Guo, et al., 2021). SA-UNet is a convolutional neural network originally designed for retinal vessel segmentation. It has two important features: a U-shaped encoder (left side)-decoder (right side) structure that achieves abstraction of spatial information while preserving the resolution, and a spatial attention unit for adaptive feature refinement (middle). The convolutional process includes $3 \times 3$ convolution, a DropBlock, a batch normalization (BN) layer, and a rectified linear unit (ReLU) (short gray right-pointing arrows). Max pooling with a stride of two handles the downsampling, then feature channels are doubled with each step (downward arrows). Decoder stages include a $2 \times 2$ transposed convolution, which halves the feature channels, and concatenation with the feature map copied from the encoder layers (upward arrows and long gray right-pointing arrows). The spatial attention module is set between the encoder and the decoder (red right-pointing arrow). In the final layer, a $1 \times 1$ convolution and sigmoid activation function is used to obtain the output segmentation map (yellow right-pointing arrow). See the original paper (Guo, et al., 2021) for further information.
(B) Raw retinal fundus vasculature image from the DRIVE dataset with a 45° field of view.
(C) Training label image with vessels denoted by white pixels. The image shown in (B) is used.
(D) A representative vasculature image of the mouse cortical surface from the training dataset. Scale bar, 1 mm.
(E) Manual annotation of the vasculature in the image shown in (D). Scale bar, 1 mm.
(F) Box chart of the time required to create one annotated image when a raw image was manually annotated (left, n = 10 images), and when a raw image was annotated by the CNN and the CNN-annotation was manually modified (right, n = 30 images).
(G) Another example input image of the mouse cortical surface for SA-UNet. This image included a very thick vessel in the midline. Scale bar, 1 mm.
(H) Example output images of vessel segmentation with SA-UNet trained with only the DRIVE dataset (left), SA-UNet trained with the DRIVE dataset and six manually created datasets (middle), and the modified SA-UNet (right). The input image is the image shown in (G). Thick vessels were incompletely segmented when the modified SA-UNet was not used (arrows). Scale bar, 1 mm.

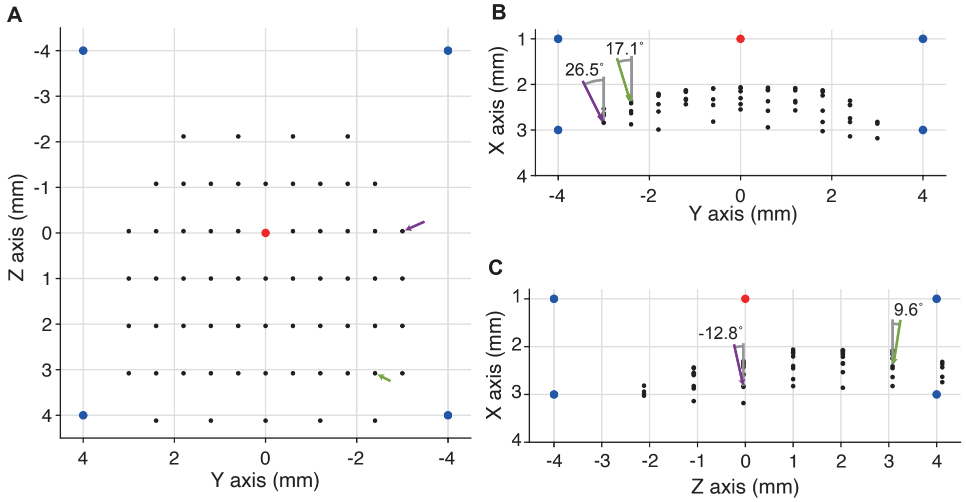


**Figure S3. Performance of TCP calibration in different coordinates.**
(A–C) Representative Y and Z positions (A), X and Y positions (B), and Z and X positions (C) of the TCP calibration points (red), calibration estimation points (blue), and the points where the cortical surface images were captured (black). Representative XYZ directions perpendicular to the cortical surface are shown by green and purple arrows. Angle values in the W and V rotations for each representative arrow are shown in (B) and (C), respectively.

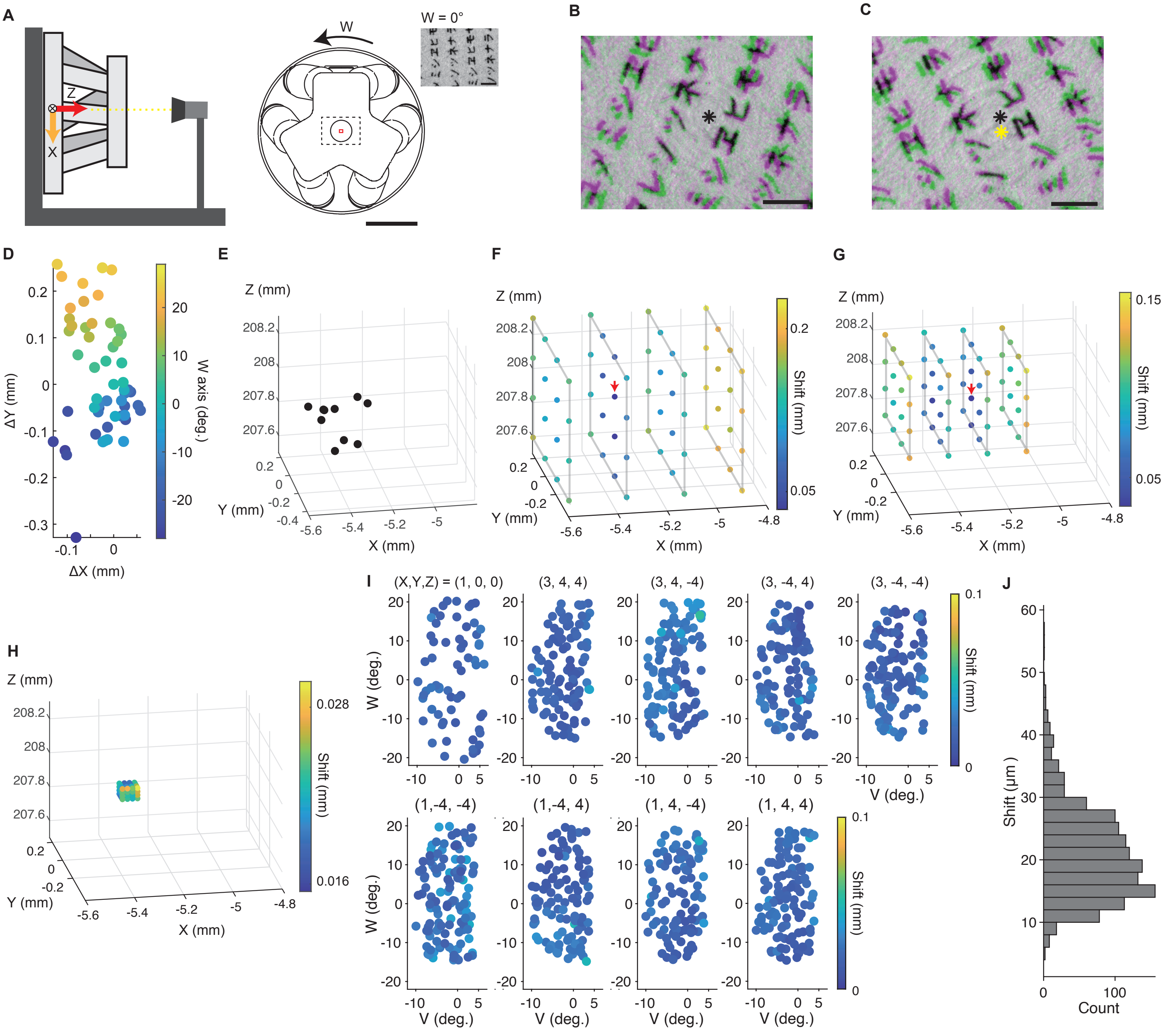


**Figure S4. Compensation for rotation error.**

(A) Schematic of the measurement of the shift by the robot stage rotation. Left: For determining the rotation center on the W-axis through multiple image captures, Camera R is positioned in front of the stage. Its optical axis is aligned along the Z axis of the base coordinate system. Right: Schematic illustration of the robot stage as viewed from the camera positioned in front of it. A paper with Japanese letters (Katakana) is affixed to the robot stage (black dashed frame). This is expanded in the inset. The red square in the diagram indicates the camera’ s field of view. Images at different W rotations were captured, as shown in (B, C). An arrow indicates the positive direction of the W rotation. Scale bar, 100 mm.

(B) Overlaid images captured at W rotations of –21° (green) and –26° (magenta). The black asterisk indicates the estimated rotation center that was calculated by comparing and matching feature points across the images. Scale bar, 1 mm.

(C) Overlaid images captured at W rotations of 26° (green) and 21° (magenta). The yellow asterisk indicates the estimated rotation center at these rotations. The black asterisk indicates the rotation center estimated from the rotations in (B). Scale bar, 1 mm.

(D) XY shift (ΔX and ΔY) of the rotation center against the angle in the W rotation. To generate each data point, an image captured at w degrees in the W rotation was compared with another captured at w + 5 degrees. The rotation center’ s coordinates were then calculated according to the procedure outlined in (A–C). These coordinates were plotted relative to their original positions at 0° in the W rotation. The pseudo-color bar indicates the degree of W rotation. n = 54 images.

(E) The coordinates of $\mathbf{P}^{E}_{Pipette}$ estimated from the pipette tip movement data, which included the XY shift of the rotation center. Each data point was calculated using measurements taken at different coordinates of the pivot point.

(F) The representative pivot points used for measuring the pipette tip shift during the first grid search. The measurement range was 0.7–0.8 mm, including all estimated coordinates in (E). A pseudo-color bar represents the extent of the pipette tip shift. A red arrow indicates the points where the shift was the smallest.

(G) The representative pivot points used for measuring the pipette tip shift during the second grid search. The measurement range was reduced to 0.47 mm, which is two-thirds of the initial range. A pseudo-color bar indicates the length of the pipette tip shift. A red arrow indicates the points where the shift was the smallest.
(H) The representative pivot points used for measuring the pipette tip shift after the final round of grid searching. The measurement range was further narrowed to 0.06 mm. A pseudo-color bar is used to illustrate the extent of the pipette tip shift.
(I) Maps of the pipette tip shift caused by rotation at different positions after applying the compensation. The coordinates of the measured positions are shown as red and blue dots in Figure S3A, B.
(J) Histogram of the pipette tip shift caused by rotations at the different positions shown in (I) (n = 1274).

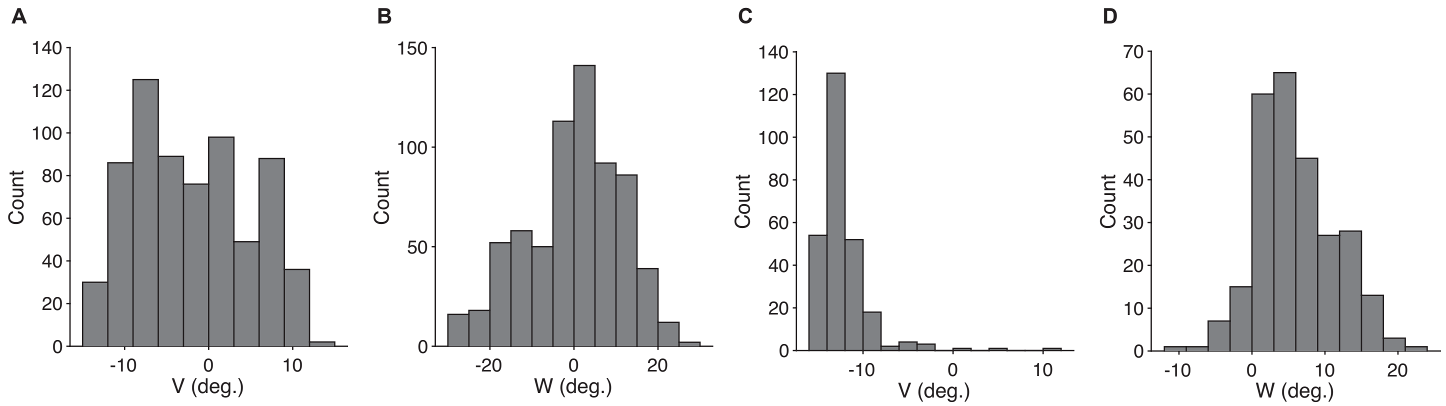


**Figure S5. Pipette insertion angles in seven mice and one marmoset.**
(A, B) Histogram of the angles in V (A) and W (B) rotations for the pipette injection into the mouse dorsal cortex (n = 679 from 7 mice).
(C, D) Histogram of the angles in V (C) and W (D) rotations for the pipette injection into the marmoset frontoparietal cortex (n = 266 from 1 marmoset).

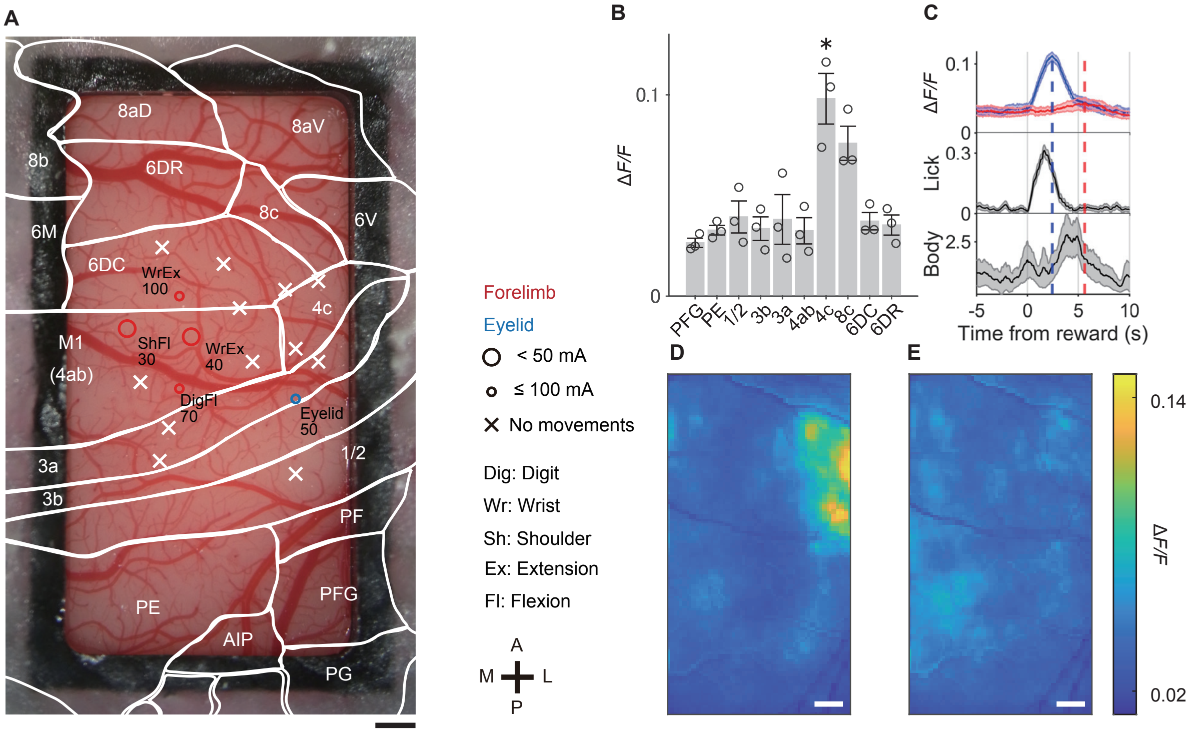


**Figure S6. Intracortical microsimulation results and fluorescence signals of different cortical areas around reward timing in the marmoset.**
(A) The results of ICMS of the sensorimotor cortex in the marmoset. Red circles indicate stimulation sites where forelimb movements were observed, and the blue circle represents a site where eyelid movement was observed. The diameters of the circles represent the minimal amplitude of the stimulation current that the movements were evoked by. Crosses indicate sites where no movements were observed. A, P, M, and L stand for anterior, posterior, medial, and lateral, respectively.
(B) Averaged Δ*F/F* values for the ten cortical areas between 1–3 s after the reward onset for individual sessions, accompanied by a bar plot representing the average value across sessions (error bar denotes the standard error). A one-way ANOVA revealed a significant difference among areas (n = 3 sessions, $F_{10,22} = 8.41$, $p = 1.6 \times 10^{-5}$). Based on the assumption that area 4ab demonstrated the strongest response to body movement, and that any cortical areas with a greater response might respond to something other than body movement, a one-sided Student' s t-test with Bonferroni correction was conducted with area 4ab against other areas to investigate whether this was true. Area 4c exhibited significantly larger responses than area 4ab. Data are plotted only for significant results using a Bonferroni correction threshold of 0.05/9 (n = 3 sessions, *p = 0.0049 < 0.0056).
(C) Δ*F/F* signals aligned to the reward delivery timing in area 4c (blue) and area 4ab (red), lick frequency, and body movements. Shading indicates ± standard error (n = 3 sessions). Dashed lines indicate the timing of the peak of Δ*F/F* signals in area 4c (blue, 2.4 s) and area 4ab (red, 5.6 s).
(D, E) Cortical maps of the averaged Δ*F/F* signals 2.4 s (D) and 5.6 s (E) after the reward delivery. Scale bar, 1 mm.

**Table S1. Experimental schedule.**

| Robot process | Time (min) | Related scripts | Animal surgery | Time (min) |
|---|---|---|---|---|
| Pipette preparation and camera focusing | 5 | main.m | | |
| Estimating rotation of the camera and the robot | 5 | TCPCalibration.m | Craniotomy of the animal | 80–100 |
| Estimating the coordinates of the needle tip | 20 | | | |
| Measuring needle dislocation due to rotation | 70 | | | |
| Scanning the brain surface | 5 | ScanSurface.m | | |
| Segmentation with SA-UNet | 1 | SA-UNet | | |
| Projecting the vessel segmentation onto the 3D surface | 5 | Autostitch, IntegrateSurface.m | | |
| Defining the area of injections manually | 1 | main.m | | |
| Calculating appropriate arrangement of injection sites | 5 | | | |
| Virus loading | 1–5 | | | |
| Automated Injection | 1 per site | | | |
| | | | Implanting glass cranial window | 5 |

**Movie S1. V-W rotation of the pipette captured by Camera L and Camera R after all calibration.**
